# Complex polyploid and hybrid species in an apomictic and sexual tropical forage grass group: genomic composition and evolution in *Urochloa* (*Brachiaria*) species

**DOI:** 10.1101/2021.02.19.431966

**Authors:** Paulina Tomaszewska, Maria S. Vorontsova, Stephen A. Renvoize, Sarah Z. Ficinski, Joseph Tohme, Trude Schwarzacher, Valheria Castiblanco, José J. de Vega, Rowan A. C. Mitchell, J. S. (Pat) Heslop-Harrison

## Abstract

**Background and Aims:** Diploid and polyploid *Urochloa* (including *Brachiaria*, *Panicum* and *Megathyrsus* species) C_4_ tropical forage grasses originating from Africa and now planted worldwide are important for food security and the environment, often being planted in marginal lands. We aimed to characterize the nature of their genomes, the repetitive DNA, and the genome composition of polyploids, leading to a model of the evolutionary pathways within the group including many apomictic species.

**Methods:** Some 362 forage grass accessions from international germplasm collections were studied, and ploidy determined using an optimized flow cytometry method. Whole-genome survey sequencing and molecular cytogenetic analysis with *in situ* hybridization to chromosomes were used to identify chromosomes and genomes in *Urochloa* accessions belonging to the different agamic complexes.

**Key Results:** Genome structures are complex and variable, with multiple ploidies and genome compositions within the species, and no clear geographical patterns. Sequence analysis of nine diploid and polyploid accessions enabled identification of abundant genome-specific repetitive DNA motifs. *In situ* hybridization with a combination of repetitive DNA and genomic DNA probes, identified evolutionary divergence and allowed us to discriminate the different genomes present in polyploids.

**Conclusions:** We suggest a new coherent nomenclature for the genomes present. We develop a model of evolution at the whole-genome level in diploid and polyploid accessions showing processes of grass evolution. We support the retention of narrow species concepts for *U. brizantha, U. decumbens*, and *U. ruziziensis*. The results and model will be valuable in making rational choices of parents for new hybrids, assist in use of the germplasm for breeding and selection of *Urochloa* with improved sustainability and agronomic potential, and will assist in measuring and conserving biodiversity in grasslands.

## Introduction

The evolution and domestication of most arable crops is well-understood and the genetic diversity of their wild relatives is being exploited in breeding new varieties. Native grasslands include high biodiversity that can be threatened by expansion of cultivated areas, while forage grasses occupy half the world’s agricultural land. Genomic knowledge is being increasingly applied to breeding the temperate *Lolium*-*Festuca* (ryegrass) complex (Velmurugan *et al*., 2016), and there are a number of genetic selection and breeding programmes for tropical and sub-tropical forage (e.g. Worthington and Miles, 2015) but applications of omics-based technologies (Ishitani *et al*., 2004) remain limited. The tropical forage grasses include clusters of species with various ploidies, and many reproduce through apomixis. Their genomic composition and diversity in general remain poorly characterized. Rational choice of parents for making crosses in breeding programmes requires knowledge of genome composition and ploidy. The integration of sequencing, molecular cytogenetic and bioinformatic tools allows the identification of genomes which come together in polyploids (Soltis *et al*., 2013). Many crop species with polyploid members, including *Brassica* (Alix *et al*., 2008) and the Brassicaceae (Cheng *et al*., 2013), *Avena* (Tomaszewska and Kosina, 2018; Liu *et al*., 2019) and particularly the tribe Triticeae (Hordeae) (Linde-Laursen *et al*., 1997) have well-established genome designations (as single letters) to describe the ancestral genomes in auto- and allo-polyploids (amphiploids). Resolution of genome relationships in the wheat group has assisted with extensive use of the germplasm pool in breeding (Feldman and Sears, 1981; Ali *et al*., 2016; Rasheed *et al*., 2018).

The pantropical grass genus *Urochloa* includes species previously classified under *Brachiaria, Megathyrsus,* and some *Eriochloa* and *Panicum* (Webster, 1987; González and Morton, 2005; Kellogg, 2015) and is a member of the Panicoideae tribe Paniceae, subtribe Melinidinae, comprising of an estimated 150 annual and perennial grasses centred in sub-Saharan Africa (Kellogg, 2015; Soreng *et al*., 2017). Joint missions in the early 1980s conducted by CGIAR (Consultative Group on International Agricultural Research) centres, CIAT (Centro Internacional de Agricultura Tropical) and ILRI (International Livestock Research Institute) in several African countries collected wild species mostly as live plant cuttings or ramets. These activities built a global grass collection with 700 accessions of *Urochloa* species representing a highly diverse gene pool for breeding and systematic studies (Keller-Grein *et al*., 1996). Valuable traits of *Urochloa* include biomass yield, physiological tolerance to low-fertility acid soils of the tropics (Arroyave *et al*., 2011), digestibility and energy content (Hanley *et al*., 2020), insect tolerance (particularly to neotropical spittlebugs; Miles *et al*., 2006) and disease resistance (Valério *et al*., 2013; Alvarez *et al*., 2014; Hernandez *et al*., 2017). However, undesirable traits are also present, such as allelopathy (leaving bare soil; Kato-Noguchi *et al*., 2014), cold-susceptibility (Mulato II; Pizarro *et al*., 2013) and invasiveness (Durigan *et al*., 2007 in the Brazilian Cerrado; *U. panicoides* is on the US Federal Noxious Weed List https://www.aphis.usda.gov/plant_health/plant_pest_info/weeds/downloads/weedlist.pdf). These *Urochloa* grass collections have huge potential for sustainable improvement as well as conservation of grasslands, including pastures, rangelands, savannah, prairie, cerrado, and roadsides and verges, with various degrees of management of grazing. Breeding or trial programmes based in Colombia, Brazil, Thailand, Zimbabwe, Ethiopia, South Africa and Australia have led to the development of over a dozen of cultivars (do Valle and Savidan, 1996; Singh *et al*., 2010) and *Urochloa* is now the most widely planted forage grass in South America occupying 60 million hectares of grasslands in the tropical savannah ecoregion of Brazil (Gracindo *et al*., 2014).

Evolution of the monophyletic *Urochloa* lineage group remains poorly understood, but it encompasses most species previously placed in the genus *Brachiaria* on morphological grounds (Webster, 1987; Salariato *et al*., 2010, 2012). The species level taxonomy within *Urochloa* established in African floras (Hutchinson and Dalziel, 1972; Clayton and Renvoize 1982; Clayton, 1989) has, however, not been fully maintained by recent floristic work (Sosef, 2016). Some *Urochloa* species have been arranged in agamic (apomictic) complexes: *U. brizantha, U. decumbens* and *U. ruziziensis* were classified into the ‘*brizantha*’ complex, and *U. humidicola* together with *U. dictyoneura* were assigned to the ‘*humidicola*’ complex (Lutts *et al*., 1991; Renvoize and Maass, 1993). Both of these species complexes have long been recognised as productive forages (Keller-Grein *et al*., 1996). Some *Urochloa* species reproduce sexually, but others with apomictic or mixed reproduction allow odd levels of ploidy and contribute to increased intraspecific variability making classification difficult. Some species are only known in the wild as diploids, but chromosome numbers of *U. ruziziensis* (Timbó *et al*., 2014) and diploid *U. brizantha* (Pinheiro *et al*., 2000) have been doubled in the laboratory to enable crossing with tetraploid apomictic species (Risso-Pascotto *et al*., 2005; de Souza-Kaneshima *et al*., 2010; Felismino *et al*., 2010). The most common basic chromosome number is *x*=9 (de Wet, 1986; Bernini and Marin-Morales, 2001), but *x*=8, *x*=7 (Basappa *et al*., 1987), and *x*=6 (Risso-Pascotto *et al*., 2006; Boldrini *et al*., 2009b; Worthington *et al*., 2019) have been reported, making the genus *Urochloa* complex.

Characterization of the genome composition and diversity of *Urochloa* germplasm, phenotypes and ploidy is required for its effective use by researchers, breeders and farmers. Both whole genome sequencing and RNA-sequencing (Higgins *et al*., in preparation) reveals unique repetitive and single-copy sequences present only in one genome and enables recognition and designation of diploid genomes and their relationships, and characterization of the genome composition in polyploids. Despite the agronomic importance of *Urochloa*, and the need to make crosses for breeding, genomes are not clearly defined (Boldrini *et al*., 2009a), although some ploidy measurements have been made (Penteado *et al*., 2000; Jungmann *et al*., 2010). The use of transposable element probes against *Urochloa* chromosomes indicates that many species are allopolyploid with differentiation in their transposable element composition (Santos *et al*., 2015), also shown by genetic analysis in apomicts (Worthington *et al*., 2016) and genomic *in situ* hybridization (Corrêa *et al*., 2020).

Here, we aimed to define the evolution and relationships of forage species in the tropical genus *Urochloa*, and understand evolutionary processes in polyploid, apomictic groups, and the diversification of abundant repetitive DNA sequences in their genomes. We measured ploidy in most of the *Urochloa* germplasm collection at CIAT (Colombia). We then aimed to use genomic and molecular cytogenetic approaches to identify repetitive DNA motifs and identify genome-specific sequences, to characterize the genomes present in the polyploid accessions (genomic composition), and develop a model of evolution at the whole-genome level in diploid and polyploid accessions in the tropical forage grass group.

## Materials and methods

### Plant material

Studies were carried out on 362 accessions of *Urochloa* and related species focusing on material available on request for research and to breeders from CIAT and USDA germplasm collections (**Supplementary Data Table S1**). We use the narrow species concepts of Clayton and Renvoize (1982) and Clayton (1989) rather than the most modern broader concepts of Sosef (2016) for ease of communication regarding the diverse genetic variants within *U. decumbens* and *U. ruziziensis*. Synonymy was updated and reconciled (POWO, 2019); many species were previously placed in *Brachiaria*, while *Urochloa maxima* is also known as *Panicum maximum* and *Megathyrsus maximus*. For accessions from the CIAT germplasm collection, RNAseq data (Higgins *et al*., in preparation) shows that 111 lines (sampled from the 362 analysed here) are genetically distinct supporting continued validity of correlation of collection locality with accession number, and a commendation to the CIAT germplasm resource collection group who maintained true lines through violent conflict, not allowing a small number of vigorous and robust lines to dominate the collection. Fresh leaf material from apomictic and sexual plants was collected in the field in Colombia and trial plots grown at CIAT and dried in silica gel. The leaf samples were then used to isolate nuclei for flow cytometry and extract whole genomic DNA. Seed samples for chromosome preparation and further cytogenetic studies were provided by Centro Internacional de Agricultura Tropical (CIAT, Colombia) and United States Department of Agriculture (USDA, USA) (**Table 1**). Voucher specimens for plants grown from seeds have been deposited in the Kew Herbarium.

**Table 1.** List of accessions used in cytological studies, their chromosome numbers and ploidy levels.

| Species | Accession number | Donor | Number of chromosomes | Base number |
| --- | --- | --- | --- | --- |
| *Urochloa* sp. | PI 657653<sup>1)</sup> | USDA | $2n=4x=32$ | 8 |
| *Urochloa* sp. | PI 508571<sup>2)</sup> | USDA | $2n=4x=36$ | 9 |
| *Urochloa* sp. | PI 508570<sup>3)</sup> | USDA | $2n=5x=45$ | 9 |
| *U. brizantha* (A.Rich.) R.D.Webster | PI 226049 | USDA | $2n=6x=54$ | 9 |
| *U. brizantha* (A.Rich.) R.D.Webster | PI 292187 | USDA | $2n=4x=36$ | 9 |
| *U. brizantha* (A.Rich.) R.D.Webster | PI 210724<sup>4)</sup> | USDA | $2n=4x=36$ | 9 |
| *U. decumbens* (Stapf) R.D.Webster | 6370 | CIAT | $2n=4x=36$ | 9 |
| *U. decumbens* (Stapf) R.D.Webster | 664 | CIAT | $2n=4x=36$ | 9 |
| *U. humidicola* (Rendle) Morrone & Zuloaga | 16867 | CIAT | $2n=8x+2/9x-4=50$ | 6 |
| *U. humidicola* (Rendle) Morrone & Zuloaga | 26151 | CIAT | $2n=6x=36$ | 6 |
| *U. ruziziensis* (R.Germ. & C.M.Evrard) | 6419 | CIAT | $2n=2x=18$ | 9 |
| *U. maxima* (Jacq.) R.D.Webster | PI 284156 | USDA | $2n=4x=32$ | 8 |
| *U. maxima* (Jacq.) R.D.Webster | 6171 | CIAT | $2n=4x=32$ | 8 |
| *U. maxima* (Jacq.) R.D.Webster | 16004 | CIAT | $2n=4x=32$ | 8 |

- 1) *Urochloa* sp. PI 657653 was received by USDA as *Panicum miliaceum*, re-identified at first as *Panicum sumatrense*, and finally described as *Urochloa* sp. We failed to determine the species identity of PI 657653, which has spikelets like *Urochloa adspersa* but leaves like *U. decumbens*, so does not match any species and may be of hybrid origin, hence the difficulties in its taxonomic identification. - 2) *Urochloa* sp. PI 508570 morphologically corresponds to *Urochloa panicoides*. - 3) *Urochloa* sp. PI 508571 was not identified. - 4) Accession PI 210724 previously assigned to *Urochloa decumbens* has been re-identified as *U. brizantha*.

### Ploidy determination

Flow cytometry was conducted to establish ploidy levels of 362 accessions of *Urochloa* and related species (355 accessions from the CIAT germplasm collection, and 7 accessions from the USDA germplasm collection; **Supplementary Data Table S1**). Cell nuclei from dehydrated leaf tissues were isolated mechanically, using method described by Doležel *et al*. (2007) with some modifications. Approximately 500 mg of tissue was chopped with razor blade in a Petri dish containing 1 mL lysis buffer (0,1M Tris, 2,5mM MgCl_2_ x 6H_2_O, 85mM NaCl, 0,1% Triton X-100; pH=7.0) supplemented with 15mM β-mercaptoethanol and 1% PVP-40 to reduce negative effect of cytosolic and phenolic compounds. The nuclear suspension was recovered and filtered through a 50-µm nylon mesh (CellTrics, Partec) to remove cell fragments and large debris, and then stained with 50 µg mL^-1^ propidium iodide, supplemented with 50 µg mL^-1^ RNase to prevent staining of double-stranded RNA. Samples were incubated on ice and analysed within 10 min in an Accuri C6 Flow Cytometer (Becton Dickinson at the Flow Cytometry Facility, University of Leicester), equipped with a 20-mW laser illumination operating at 488 nm. The results were acquired using the CFlow® Plus software. The software was set up according to Galbraith and Lambert (2012). Ploidy levels of *Urochloa* were estimated by comparing the PI fluorescence intensities of samples of target species to that of samples of an internal standard (*U. maxima* CIAT 6171 2*n*=4*x*=32, *U. ruziziensis* CIAT 26433 2*n*=2*x*=18 or *U. humidicola* CIAT 26151 2*n*=6*x*=36). The coefficient of variation (CV) of the G_0_/G_1_ peak was evaluated in each sample to estimate nuclei integrity and variation in DNA staining.

### DNA extraction and sequencing

Genomic DNA was extracted from fresh and dried leaves with the use of standard cetyltrimethylammonium bromide (CTAB)-based method (Doyle and Doyle, 1990) with minor modifications. Whole genomic DNA from nine *Urochloa* accessions of various ploidies (**Supplementary Data Table S2)** was sequenced commercially (Novogene) with Illumina HiSeq 2x 150bp paired-end reads (c. 12Gbases). Project data have been deposited at the National Center for Biotechnology Information (NCBI; https://www.ncbi.nlm.nih.gov/sra/) under BioProject …………………. (SRR………………….).

**Table 2.** Number of analyzed accessions and their distribution in the various levels of ploidy.

| Species | $2n=2x$ | $2n=4x$ | $2n=5x$ | $2n=6x$ | $2n=7x$ | $2n=8x+2$<br>or<br>$2n=9x-4$ | $2n=9x$ |
| --- | --- | --- | --- | --- | --- | --- | --- |
| *Andropogon gayanus* | 1 |  |  |  |  |  |  |
| *Panicum coloratum* | 1 |  |  |  |  |  |  |
| *Paspalum dilatatum* |  | 1 |  |  |  |  |  |
| *Pennisetum polystachion* |  |  |  | 1 |  |  |  |
| *Pennisetum purpureum* |  | 1 |  |  |  |  |  |
| *Urochloa arrecta* |  | 1 |  |  |  |  |  |
| *Urochloa brizantha* | 6 | 59 | 25 | 2 |  |  |  |
| *Urochloa decumbens* | 18 | 26 |  | 1 |  |  |  |
| *Urochloa dictyoneura* |  |  |  |  | 1 |  |  |
| *Urochloa dura* |  |  |  | 1 |  |  |  |
| *Urochloa humidicola* |  |  |  | 16 | 33 | 1 | 3 |
| *Urochloa jubata* | 1 | 1 |  |  |  |  |  |
| *Urochloa maxima* | 25 | 104 |  |  |  |  |  |
| *Urochloa nigropedata* |  | 1 |  |  |  |  |  |
| *Urochloa plantaginea* | 1 |  |  |  |  |  |  |
| *Urochloa platynota* | 1 |  |  |  |  |  |  |
| *Urochloa ruziziensis* | 26 |  |  |  |  |  |  |
| *Urochloa hybrid* |  | 1 |  |  |  |  |  |
| *Urochloa* sp. 1 |  |  | 1 |  |  |  |  |
| *Urochloa* sp. 2 |  | 1 |  |  |  |  |  |
| *Urochloa* sp. 3 |  | 1 |  |  |  |  |  |

### Analyses of genomic abundance

Whole genome sequencing data were used to discover most abundant repeats, and establish genomic compositions of *Urochloa* accessions of different ploidy levels. Whole genome shotgun sequence from *U. ruziziensis* cultivar CIAT 26162 (deposited in SRA under accession PRJNA437375; Worthington *et al*., 2021) was used as a reference genome. Highly abundant repetitive DNA sequences were extracted as high-frequency 50-mers using the program Jellyfish v.2.2.6 (Marçais and Kingsford, 2011). Similarity-based clustering, repeat identification, and classification of a subset raw reads were performed using RepeatExplorer (Novak *et al*., 2013) and TAREAN (Novak *et al*., 2017). All potential specific sequences extracted as 50-mer repeats or clusters were mapped to reference genome and paired reads from nine sequenced genomes using the program Geneious (Kearse *et al*., 2012). 50-mer repeats, contigs and clusters were analysed by BLAST search against NCBI database to check for repeat identification (Sayers *et al*., 2019). The PCR primer pairs were designed using the Primer3 (Rozen and Skaletsky, 1999).

### Probes used for in situ hybridization

Four different types of probes were used for FISH:

1. two ribosomal DNA sequences: pTa71 (Gerlach and Bedbrook, 1979) which contains a 9-kb EcoRI fragment of *Triticum aestivum* L. consisting of the 18S-5.8S-25S rRNA genes and the transcribed and non-transcribed intergenic spacer regions; pTa794 (Gerlach and Dyer, 1980) which contains part of the *T. aestivum* 5S rRNA gene and spacer sequences.
2. whole genomic DNA extracted from six diploid species (**Supplementary Data Table S11**).
3. conserved regions found by k-mer and RepeatExplorer analysis, and amplified using newly designed genome-specific primers (**Supplementary Data Table S3; Supplementary Data Table S9**)
4. newly designed synthetic 50-mer oligonucleotide probes (**Supplementary Data Table S3**).

Probes 1-3 were labelled with digoxigenin-11-dUTP or biotin-11-dUTP (Roche) using BioPrime® Array CGH, and then purified using BioPrime® Purification Module (Invitrogen). Fluorescent nucleotides were incorporated during synthesis for probes from group 4.

### Chromosome preparation

Chromosome preparation was carried out using the method previously described by Schwarzacher and Heslop-Harrison (2000). The root-tips were collected from plants cultivated in a greenhouse, treated with α-bromonaphthalene at 4°C for 6 h to accumulate metaphases, and fixed in 3:1 ethanol:acetic acid. Fixed root-tips were washed in enzyme buffer (10mM citric acid/sodium citrate) for 15 min, digested in enzyme solution: 20U/ml cellulase (e.g. Sigma C1184), 10U/ml ‘Onozuka’ RS cellulase and 20U/ml pectinase (e.g. Sigma P4716 from *Aspergillus niger*; solution in 40% glycerol) in 10mM enzyme buffer, and squashed in 60% acetic acid. Cover slips were removed after freezing with dry ice. Slides were air-dried and used for *in situ* hybridization within 3 months.

### In situ hybridization procedure

Fluorescent *in situ* hybridization was carried out using the method described by Schwarzacher and Heslop-Harrison (2000), with minor modifications. The hybridization mixture and the slides were denatured together in a hybridization oven for 7 min at 75°C. Hybridization was performed at 37°C overnight or 2 days depending on the type of probe. Post-hybridization washes were carried out at 42°C: in 2x SSC for 2 min, in 0.1x SSC or 20% formamide in 0.1x SSC for 6 min (depending on the type of probe), and 2x SSC for 20 min. Hybridization sites were detected with streptavidin conjugated to Alexa 594 (Life Technologies-Molecular Probes) and antidigoxigenin conjugated to fluorescein isothiocyanate (FITC) (Roche Diagnostics). Slides were then counterstained with DAPI. Mounted slides were examined with a Nikon Eclipse 80i epifluorescence microscope, and photographs were taken using a DS-QiMc monochromatic camera, and NIS-Elements v.2.34 software (Nikon, Tokyo, Japan) and assembled in Photoshop (Adobe) using only software functions affecting the whole image.

## Results

### Taxonomic identification and ploidy measurement

We studied 362 accessions of *Urochloa* and related genera (17 species, 1 synthetic hybrid, 3 unidentified accessions; **Table 1; Table 2; Supplementary Data Table S1**), and verified these taxonomically using live plants in CIAT, Cali, Colombia and reference herbarium specimens at the Royal Botanic Gardens, Kew, UK, linked where available with collection localities in Africa, morphological traits and cultivar status (CIAT Genebank data, Genesys database, and unpublished reports). In some cases, species identity had to be revised. Ploidy levels were determined using flow cytometry of fluorescently stained nuclei from dried leaf materials with a newly optimized method achieving a CV (coefficient of variation) of typically 2-5% (**Supplementary Data Fig. S1**); this presents a marked improvement for dried material over the previous method of Doležel *et al*. (2007).

Ploidy levels of 2*x*, 4*x*, 5*x*, 6*x,* 7*x,* and 9*x* were found (**Table 2**), but some accessions differed from published values (**Supplementary Data Table S1**). *U. ruziziensis* was only found as a diploid (2*n*=2*x*=18), while other species, e.g. *U. humidicola,* were found only as polyploids (6*x*, 7*x*, and 9*x*). No correlation between the level of ploidy of the examined accessions of *U. brizantha*, *U. decumbens, U. ruziziensis,* and *U. humidicola*, with the area of their original East African collection sites was evident as shown on the maps of **Fig. 1**. A mixture of species and ploidy levels was found at most collection sites; 4*x* and 5*x* accessions were predominant for *U. brizantha*, and 6*x* and 7*x* for *U. humidicola* (see also **Table 2**).

**FIG. 1.**
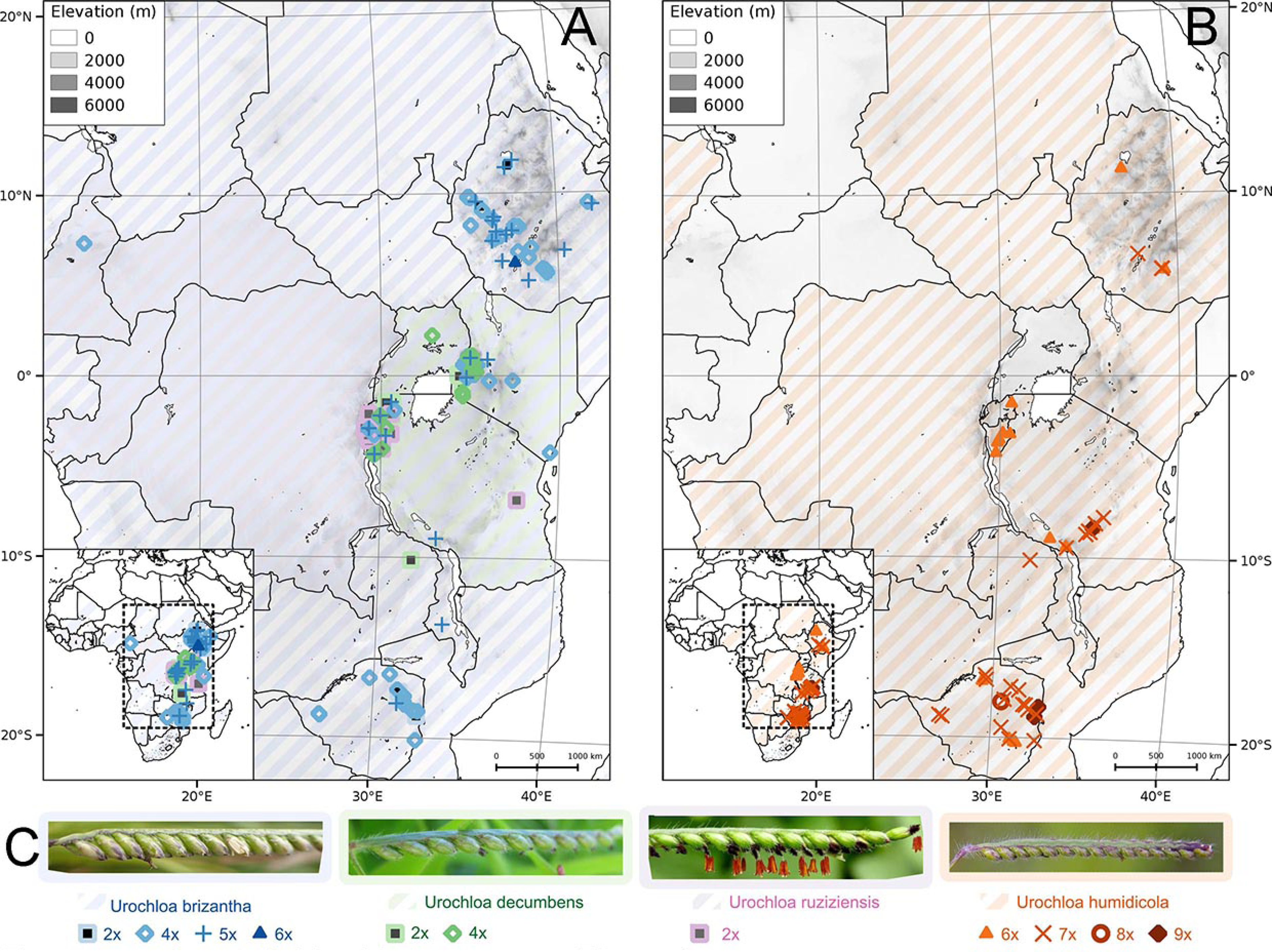
The natural distribution ranges of *Urochloa* (diagonal shading) and geographical origin of accessions studied here (symbols). For relevant species, multiple ploidies were found at each collection location. (A) *U. brizantha* is marked in blue, *U. decumbens* in green, *U. ruziziensis* in pink. (B) *U. humidicola* is marked in orange. (C) Spikelet morphology. Diploid accessions are shown as squares, tetraploid as empty diamonds, pentaploid as upright crosses, hexaploid as triangles, heptaploid as diagonal crosses, octoploid as circles, and nonaploid as filled diamonds. Natural distribution ranges are from wcsp.science.kew.org. Photograph of *U. humidicola* by Russell Cumming.

### Number of chromosomes and rDNA sites

Studied accessions were euploid with basic chromosome numbers of *x*=6 for *U. humidicola*; *x*=8 for *U. maxima*; and *x*=9 for *U. brizantha*, *U. decumbens* and *U. ruziziensis* (**Table 1**) with the exception of one accession of *U. humidicola* CIAT 16867 with 2*n*=50 (2*n*=8*x*+2 or 9*x*-4). Fluorescent *in situ* hybridization with a wheat 45S rDNA (18S-5.8S-25S; probe pTa71) and 5S rDNA (probe pTa794) (**Fig. 2**) showed, typically, one pair of major 45S rDNA sites per genome, and two pairs of 5S sites on different chromosomes in species belonging to *’brizantha’* complex. In *U. maxima*, one pair of 45S rDNA sites and one pair of 5S rDNA sites per genome were observed. *U. humidicola* had one pair of chromosomes showing both 45S and 5S rDNA signals. The pattern of rDNA sites in unidentified accessions PI 657653 and PI 508570 did not correspond to other tetraploids studied here.

**FIG. 2.**
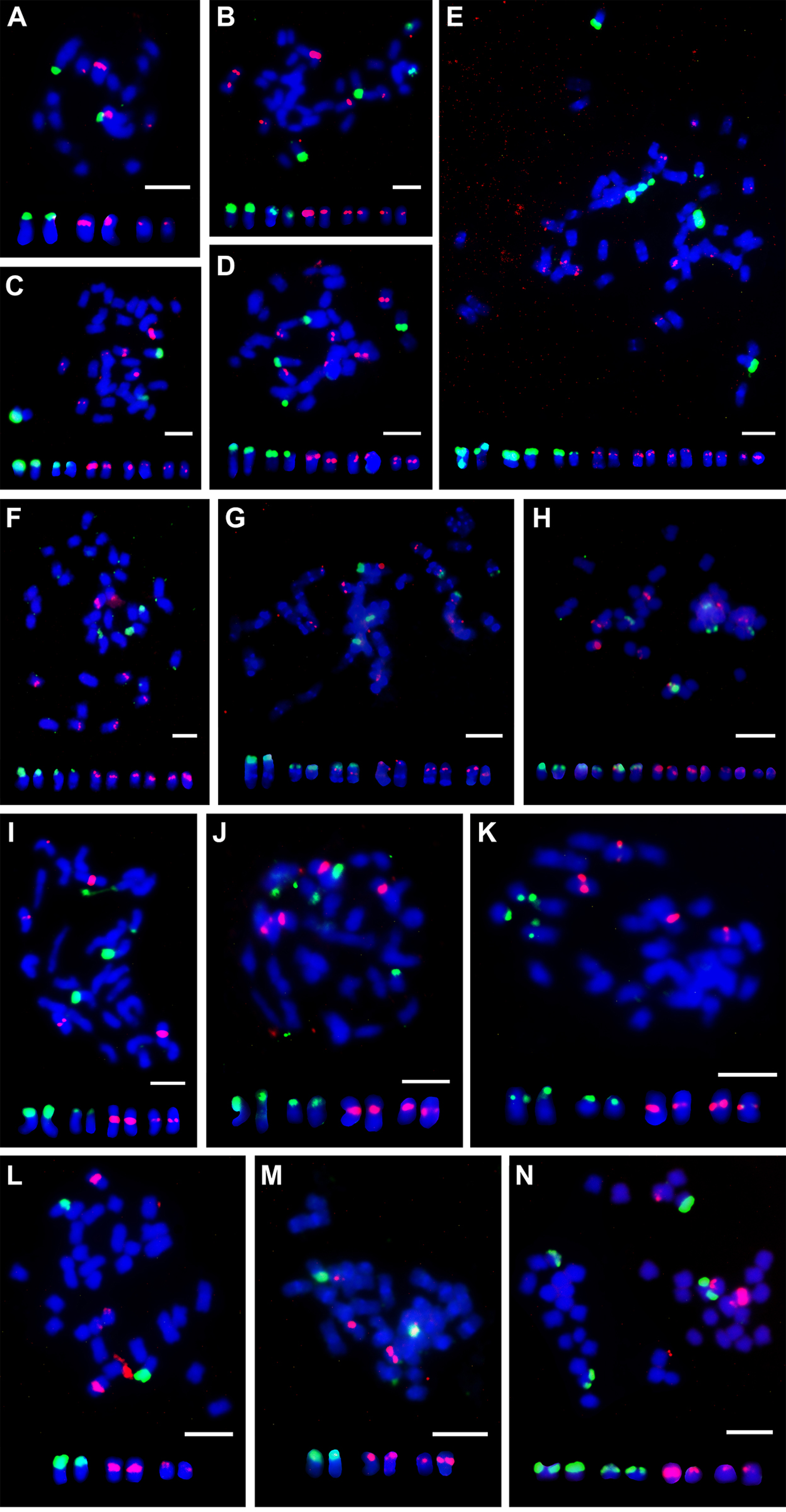
Localization of ribosomal 5S (red) and 18S-5.8S-25S (green) DNA sites on metaphase chromosomes of *Urochloa* species by fluorescent *in situ* hybridization. (A) *U. ruziziensis* (2*x*; CIAT 6419), one pair of chromosomes with large 18S-5.8S-25S locus detected on the satellites, and two pairs of chromosomes with 5S loci observed in the interstitial regions. (B) *U. decumbens* (4*x*; CIAT 664), 18S-5.8S-25S locus on satellites of two chromosome pairs, and 5S observed in the interstitial regions of three chromosome pairs; one pair of chromosomes showed much stronger 5S rDNA signals, as did diploid *U. ruziziensis*. (C) *U. decumbens* (4*x*; CIAT 6370), same number and position of rDNA signals as in *U. decumbens* CIAT 664 (B). (D) *U. brizantha* (4*x*; PI 210724), same number and position of rDNA signals as in tetraploid *U. decumbens* CIAT 664 (B) and CIAT 6370 (C). (E) *U. brizantha* (6*x*; PI 226049), three pairs of chromosomes with 18S-5.8S-25S rDNA sites and five pairs of chromosomes with 5S rDNA sites. (F) *U. brizantha* (4*x*; PI 292187), same number and position of rDNA signals as in tetraploid *U. decumbens* CIAT 664 (B) and CIAT 6370 (C). (G) *U. humidicola* (6*x*; CIAT 26151), three pairs of chromosomes showed 18S-5.8S-25S rDNA signals on their satellites, and one pair of them was different, having additional 5S rDNA sites; another three pairs of chromosomes had 5S rDNA signals. (H) *U. humidicola* (8*x*+2 or 9*x*-4; CIAT 16867), three pairs of chromosomes showed 18S-5.8S-25S rDNA signals, and one pair of them had additional 5S rDNA sites; another four pairs of chromosomes had 5S rDNA signals. (I) *U. maxima* (4*x*; CIAT 6171), two pairs of chromosomes with large 18S-5.8S-25S rDNA signals, and two pairs of chromosomes with 5S rDNA signals detected at the centromeric-pericentromeric regions. (J) *U. maxima* (4*x*; CIAT 16004), same number and position of rDNA signals as in *U. maxima* CIAT 6171 (I). (K) *U. maxima* (4*x*; PI284156), same number and position of rDNA signals as in CIAT 6171 (I) and CIAT 16004 (J). (L) *Urochloa* sp. (4*x*; PI 657653), one pair of chromosomes with 18S-5.8S-25S rDNA signals and two pairs of chromosomes with 5S rDNA, which does not correspond to the pattern of rDNA signals in other tetraploids studied here. (M) *Urochloa* sp. (4*x*; PI 508571), same number of rDNA signals as in PI 657653 (L). (N) *Urochloa* sp. (5*x*; PI 508570), two pairs of chromosomes showing 18S-5.8S-25S rDNA and two pairs of chromosomes with 5S rDNA signals. Scale bars = 5µm

### Repetitive DNA motifs identified by k-mer and graph-based clustering of DNA sequence reads

The most abundant 50-mer repeats were extracted from 2Gb of whole genome sequence reads from each of the ten accessions (our whole genome sequencing data from nine accessions along with published whole genome shotgun sequence from *U. ruziziensis*, Worthington *et al*., 2021). Those with sequence homology to rDNA, sequencing primers or chloroplast genomes, or with extreme GC ratio were omitted from further analysis as well as the telomeric repeat, (TTTAGGG)_n_, that represented about 0.1% of the reads. The genome proportion of remaining abundant 50-mers from reads or from assemblies of abundant 50-mers into contigs showed most, but not all, motifs were on the whole present with similar abundance in most *Urochloa* accessions (**Supplementary Data Table S3;** BLASTN search in **Supplementary Data Table S4)** indicating only limited species or genome-specific repeats are present. Comparison of abundant 50-mer motifs from the three diploid species, *U. ruziziensis* CIAT 26162, *U. decumbens* CIAT 26305 and *U. maxima* CIAT 16049, did not reveal any *U. decumbens*-specific repeats. But some repeats showed different abundance: repeat 1392_24 was much more abundant in *U. ruziziensis* (1.18%, **Supplementary Data Table S3**), repeat 1101_42 represents 0,43% of the *U. maxima* genome, and has very low homology to *U. brizantha* and *U. humidicola* accessions. Repeat 1162_31 represents 1.72% of the *U. maxima* genome, and is also highly abundant in genomes of two polyploid accessions of *U. brizantha*, representing more than 1% of their genomes.

Since the whole genome sequence of diploid *U. brizantha* was not available, two k-mer strategies were used to find sequences potentially specific for the *U. brizantha* genome. In the first, the abundant 50-mer motifs from four polyploid accessions of *U. brizantha* were mapped to each other. Sequences that occurred in all four accessions were then *de novo* assembled. Contig 5 (**Supplementary Data Fig. S2**) with the highest genome proportion in *U. brizantha* accessions, but no or very low genome proportion in diploid *U. decumbens* and *U. ruziziensis,* was a candidate motif specific to the genome of *U brizantha*. In the second strategy, we tested our hypothesis (built on initial FISH results) that tetraploid *U. decumbens* is an allopolyploid with the genomic composition XXYZ (where X, Y and Z represent genomes to be determined). We mapped abundant unassembled 50-mer motifs from tetraploid *U. decumbens* CIAT 664, to 50-mer datasets from diploid *U. decumbens*, *U. ruziziensis* and *U. maxima*. The differentially-abundant sequence 1771_76 (high genome proportion in tetraploid *U. decumbens* and four *U. brizantha* accessions but low in the diploids; **Supplementary Data Table S3**) was a candidate repeat specific to genome Z.

The 50-mer sequence dataset from *U. humidicola* (6*x*) was mapped to highly abundant 50-mers from the three diploid species and identified four motifs unique to *U. humidicola*: 5899, 1014, 1015_2, and 7000 (**Supplementary Data Table S3**). Similarly, to find abundant repetitive motifs in 4*x Urochloa* sp. (PI 657653, unidentified hybrid), two 50-mer sequences were extracted, 1134_5 and 1644_4, representing 0.33% and 0.2% of the genome, respectively, with very low abundance in the diploids.

For graph-based sequence clustering and characterization of repeats, 2Gb subsets of raw sequence from each of the nine *Urochloa* accessions were analysed using RepeatExplorer and TAREAN (Novak *et al*., 2013; Novak *et al*., 2017). Generally, 38.2% to 60.0% of reads were assigned into clusters of related sequence reads (**Supplementary Data Fig. S3**; **Supplementary Data Table S5; Supplementary Data Table S6; Supplementary Data Table S7; Supplementary Data Table S8**). As with k-mers, sequences showing high homology to rDNA or chloroplast genomes, and extreme GC ratio were omitted from further analysis, and the final list of putative genome-specific sequence was created by comparing genome abundance between accessions (**Supplementary Data Table S9**). The number of raw reads with high homology to the most abundant clusters/RE motifs in each one of the sequence datasets were then counted for ten whole genome sequence reads. Those clusters showing high proportion in one diploid genome are candidate genome-specific sequences (**Supplementary Data Table S9**), and some were selected for testing as probes by chromosomal *in situ* hybridization (see below).

Transposable elements were recognized in each of the nine sequenced genomes (**Fig. 3; Supplementary Data Table S10**; for *U. ruziziensis* see Worthington *et al*., 2021). The automated annotation provided by RepeatExplorer will omit, or give wrong, identification of some elements based on homology to known sequences; therefore, regardless of annotation, any elements differing in abundance between accessions were candidates for use as probes for *in situ* hybridization to distinguish genomes. Thus, sequences with differential abundance identified were the Bianca retrotransposon in *U. brizantha* polyploids, the highly abundant Takey retrotransposon in diploid *U. decumbens*, the retrotransposon CRM in 4x PI 657653, and the LINE in *U. humidicola* (arrows in **Fig. 3**). The Tork retrotransposon was found in some *U. brizantha*, indicating the differences in genome structure between accessions.

**FIG. 3.**
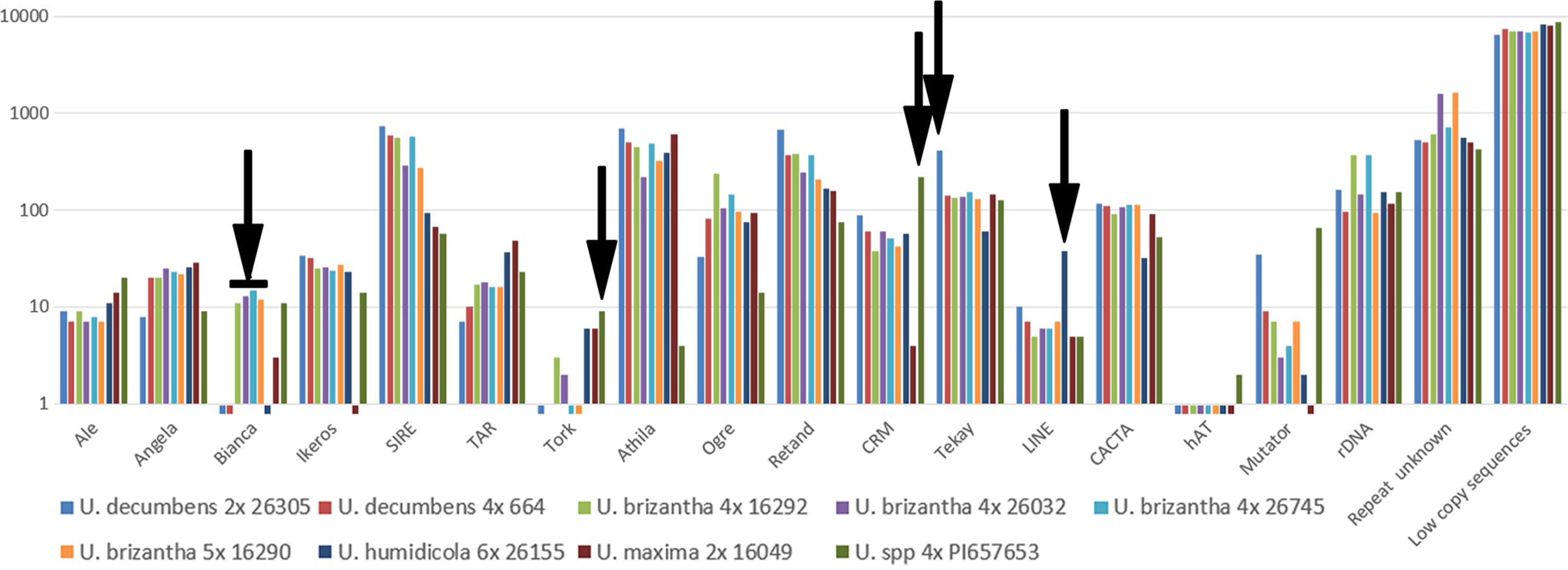
Relative abundance (log) of DNA sequence classes in *Urochloa* species and hybrids from whole-genome sequence reads. Automated repeat identification of graph-based clustering (RepeatExplorer, TAREAN) and abundant k-mer motifs, were classified by nucleotide domain hits and database BLASTN searches (Supplementary Data Table S6; Supplementary Data Table S8). Arrows indicate some motifs with differential abundance between accessions. Bars below 1 indicate abundance <0.01% of genome.

### Organization and genome-specificity of repetitive DNA sequences on chromosomes

Total genomic DNAs (gDNA) from diploid species of *U. brizantha* (Ubriz), *U. decumbens* (Udec), *U. ruziziensis* (Uruz) and *U. maxima* (Umax) were used as probes for genomic *in situ* hybridization (**Supplementary Data Table S11**) on 14 accessions of *Urochloa* diploids *and* polyploids. Results are described in **Supplementary Data Table S12**, and example micrographs are shown in **Fig. 4** (giving details about probe combinations and observed signals in the legend). In summary, gDNA_Uruz and gDNA_Udec probes showed signals in centromeric-pericentromeric regions rather than painting whole chromosomes, and the differential hybridization strengths (hybridization stringency 72% and 85%) did not affect GISH results. Different intensities of centromeric-pericentromeric signals in polyploids belonging to ‘*brizantha*’ complex indicate that these species are allopolyploids (**Figs. 4A-J**). The simultaneous use of gDNA_Uruz and gDNA_Udec probes against chromosomes of *U. humidicola* showed differential dispersed signals on many chromosomes, indicating the differences between diploid *U. ruziziensis* and *U. decumbens* genomes (**Fig. 4K**). gDNA_Ubriz probe hybridized to rDNA sites of different species, but not to centromeric-pericentromeric regions of chromosomes (**Fig. 4I; Supplementary Data Table S12**). gDNA_Umax probe showed very strong centromeric-pericentromeric signals on all 32 chromosomes of tetraploid *U*. maxima, and addition to terminal and subterminal regions (**Fig. 4L** and **Fig. 7**). *Urochloa* accessions not assigned to species are clearly allopolyploids (**Fig.** 4N, O; Supplementary Data Table S12).

**FIG. 4.**
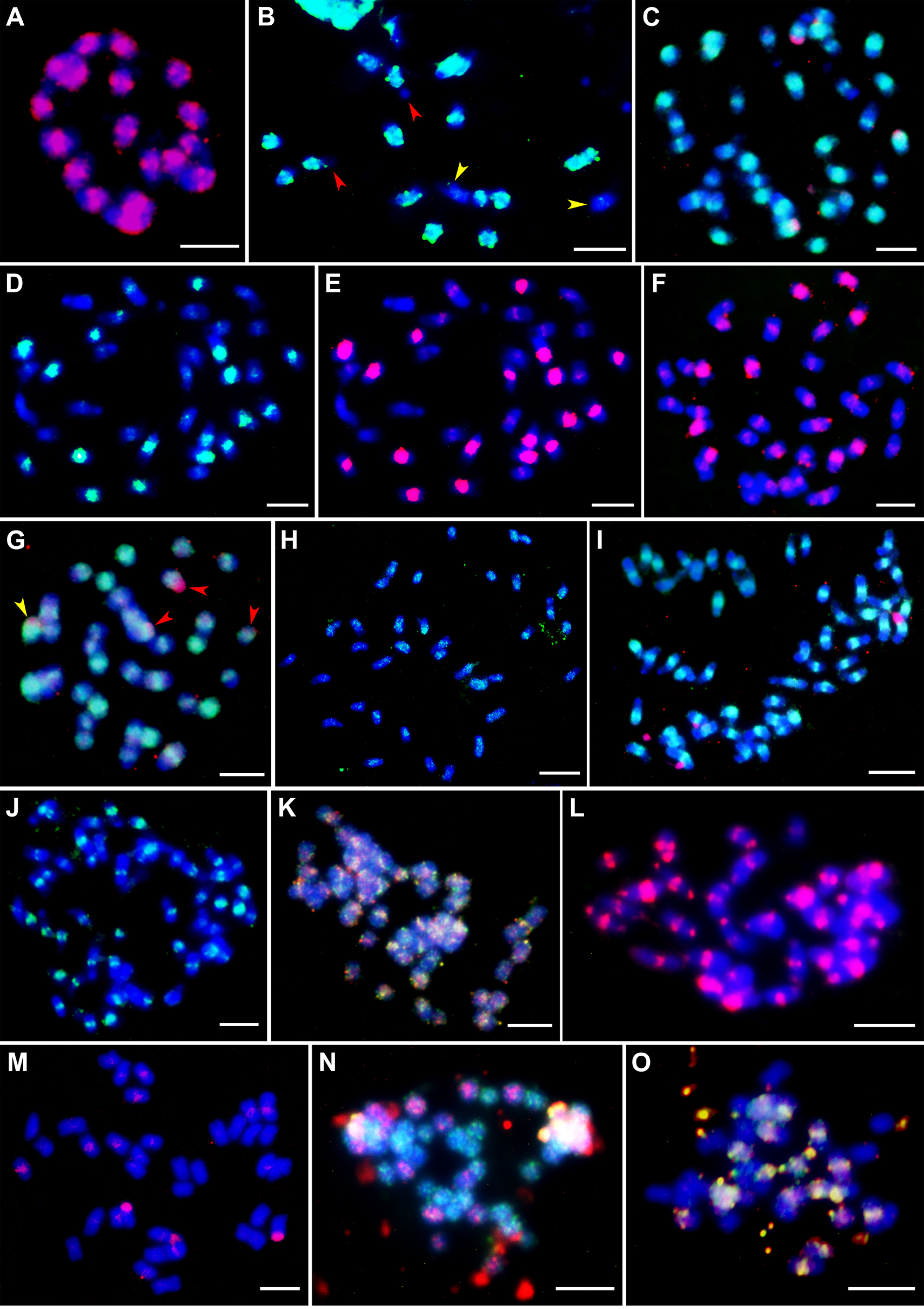
Localization of labelled whole genomic DNA (gDNA) from diploid species used as probes for *in situ* hybridization on metaphase chromosomes (fluorescing blue). (A) 18 chromosomes of *U. ruziziensis* (2*x*, CIAT 6419) showing strong signals of gDNA_Uruz1 probe (red) in centromeric-pericentromeric regions of chromosomes and some terminal signals on one or both arms of several chromosomes; (B) *U. ruziziensis* (2*x*, CIAT 6419) metaphase showing strong signals of gDNA_Udec1 probe (green) in 8 pairs of chromosomes (satellites of two chromosomes remain unlabelled; red arrowheads), and weak signals in centromeres of 2 chromosomes (yellow arrowheads); (C) metaphase of *U. decumbens* (4*x*, CIAT 6370) with 18 strong and 18 weak signals of gDNA_Udec2 probe (green), and 4 red signals of rDNA after hybridization with gDNA_Umax1 probe; (D) metaphase of *U. decumbens* (4*x*, CIAT 6370) showing 18 strong and 18 weak signals of gDNA_Udec1 probe (green); (E) same metaphase as in D, showing signals of gDNA_Uruz1 probe (red) in the same position as gDNA_Udec1 probe, which confirms the similarity of the genomes of these accessions; (F) chromosomes of *U. brizantha* (4*x*, PI 292187) with red signals of gDNA_Uruz1 probe: 9 very strong and 27 weak; (G) chromosomes of *U. brizantha* (4*x*, PI 292187) with green signals of gDNA_Udec1 probe: 9 very strong and 27 weak; gDNA_Umax1 probe shows 4 red signals (rDNA): 1 signal in chromosome with strong gDNA_Udec1 signal (yellow arrowheads) and 3 signals in chromosomes with weak gDNA_Udec1 signals (red arrowheads); (H) metaphase of *U. brizantha* (4*x*, PI 292187) with some rDNA and dispersed signals of gDNA_Umax2 probe (green); gDNA_Ubriz1 probe does not show signals; (I) metaphase of *U. brizantha* (6*x*, PI 226049) with gDNA_Uruz1 probe signals (green) in centromeres: 18 strong, 18 weaker and 18 weak; gDNA_Ubriz1 probe (red) gives 4 red signals (rDNA); (J) metaphase of *U. brizantha* (6*x*, PI 226049) with gDNA_Udec1 probe signals (green) in centromeres: 18 strong, 18 weaker and 18 weak; (K) metaphase of *U. humidicola* (8*x*+2 or 9*x*-4, CIAT 16867) showing dispersed signals of gDNA_Uruz1 (red) and gDNA_Udec2 (green) probes along chromosomes; many of these signals do not overlap; (L) metaphase of *U. maxima* (4*x*, CIAT 6171) with centromeric and telomeric signals of gDNA_Umax1 probe (red); (M) chromosomes of *U. decumbens* (4*x*, CIAT 664) with rDNA and very weak dispersed signals in centromeres after hybridization with gDNA_Umax1 probe (red); (N) metaphase of *Urochloa* sp. (5*x*, PI 508570) showing 18 chromosomes with dispersed signals of gDNA_Ubriz1 probe (red) and 27 chromosomes with dispersed signals of gDNA_Umax1 probe (green); (O) metaphase of *Urochloa* sp. (4*x*, PI 508571) showing signals of gDNA_Uruz1 probe (red) and gDNA_Udec1 probe (green) in 18 chromosomes; the 18 remaining chromosomes show no signals. Scale bars = 5µm

Probes designed from highly abundant sequences recognized by k-mer (**Supplementary Data Table S3**), and RepeatExplorer and TAREAN (**Supplementary Data Table S9**) analyses, were used, mostly in differential pairs, for *in situ* hybridization to localise repeats on *Urochloa* chromosomes, and distinguish genomes in polyploids (**Fig. 5** for *’brizantha*’ and **Fig. 6** for ‘*humidicola*’ complexes; signal summary in **Supplementary Data Table S3** and **Supplementary Data Table S9**; chromosomes were grouped by signal location and intensity). Overall, *in situ* hybridization strength correlated with *in silico* analysis (percentage of sequence in the genomes), now showing genome and chromosomal distribution of the probes and enabling discrimination of the genome of origin of most chromosomes in the polyploid accessions. All putative Ubriz-specific probes designed using two strategies gave the same number and position of signals; and some chromosomes shared both Ubriz- and Udec-specific signals. Detailed descriptions of the probes and hybridization results are given in the extended legend (**Figs. 5** and **6**) and probe description in **Supplementary Data Table S3** and **Supplementary Data Table S9**.

**FIG. 5.**
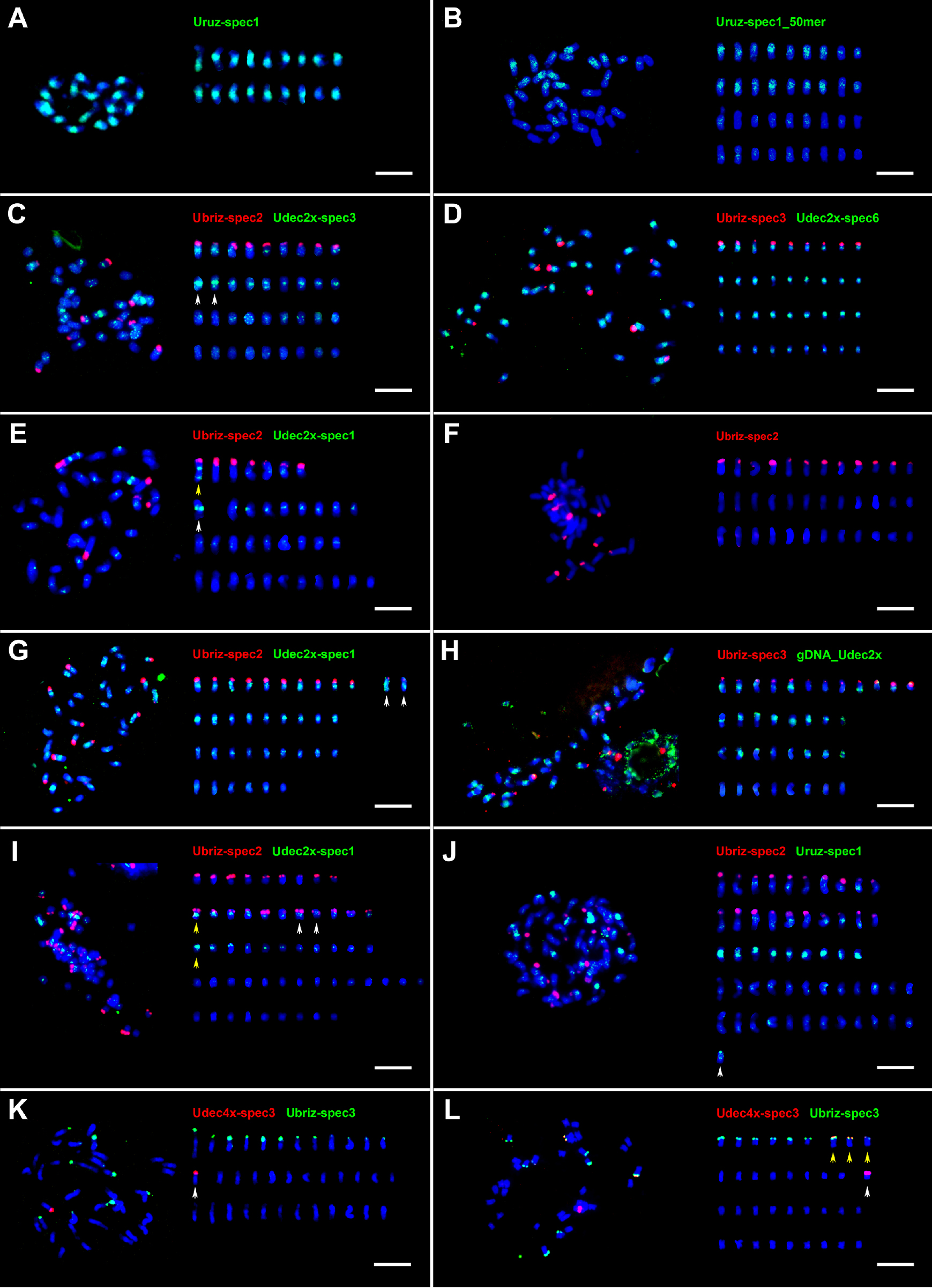
Localization of abundant repetitive sequences on chromosomes of different species belonging to the ‘*brizantha*’ complex. Probes are described in Table S3. Chromosomes (right) were arranged by size and FISH signal. (A) 18 strong signals of Uruz-spec1 probe (green) at centromeric-pericentromeric regions of *U. ruziziensis* (2*x*, CIAT 6419) chromosomes; (B) Uruz-spec1_50mer labelled 18 chromosomes of *U. decumbens* (4*x*, CIAT 664; green); (C) Ubriz-spec2 probe showed strong signals in terminal regions of nine chromosomes of *U. decumbens* (4*x*, CIAT 664). Udec2x-spec3 showed strong signals in centromeric-pericentromeric positions of chromosomes, even those with terminal Ubriz-spec2 signals. Two chromosomes exhibit very strong green fluorescence (white arrowheads). Some signals were more dispersed along chromosomes, and the 18 chromosomes without Ubriz-spec2 signal had weak Udec2x-spec3 signals; (D) metaphase of *U. decumbens* (4*x*, CIAT 664); Ubriz-spec3 showed a similar pattern of signals as Ubriz-spec2 in C. Udec2x-spec6 showed only 9 chromosomes with weak signals; (E) metaphase of *U. decumbens* (4*x*, CIAT 6370); Ubriz-spec2 probe produced seven signals in the terminal position of chromosomes; one chromosome with (yellow arrowhead) and one chromosome without Ubriz-spec2 signals (white arrowhead) showed strong signals in centromeric and subtelomeric positions of Udec2x-spec1 probe; (F) metaphase of *U. brizantha* (4*x*, PI 210724); 12 signals of Ubriz-spec2 probe at terminal regions of chromosomes; (G) metaphase of *U. brizantha* (4*x*, PI 292187); same number and position of Ubriz-spec2 signals as in F where 2 of the 12 signals were weaker. Thirty chromosomes showed strong- to weak-Udec2x-spec1 signals, while the other six had very weak or no signals; (H) *U. brizantha* (4*x*, PI 292187); gDNA-Udec probe gave strong signals on some chromosomes with Ubriz-spec3 signals and those without Ubriz-spec3 signals; (I) *U. brizantha* (6*x*, PI 226049); Ubriz-spec2 and Udec2x-spec1 probes differentiate chromosomes into five types: 9 chromosomes with Ubriz-spec2 signals (group I), 11 chromosomes with Ubriz-spec2 and Udec2x-spec1 signals (group II), 11 chromosomes with strong Udec2x-spec1 signals (group III), 14 chromosomes with very weak Udec2x-spec1 signals (group IV), 9 chromosomes without any signals (group V). In the group II, there was a pair of chromosomes showing the same pattern of signals (white arrowheads), although it seems that another chromosome from this group (yellow arrowhead) had the same strong centromeric-pericentromeric signal of Udec2x-spec1 probe as another chromosome from the group III (yellow arrowhead); (J) metaphase of *U. brizantha* (6*x*, PI 226049) showing 9 chromosomes with very strong centromeric-pericentromeric signals of Uruz-spec1 probe; (K) metaphase of *U. brizantha* (4*x*, PI 292187); Ubriz-spec3 probe gave 12 signals at terminal position of chromosomes. One very strong signal of Udec4x-spec3 detected at terminal region of one chromosome; (L) metaphase of *U. decumbens* (4*x*, CIAT 664) showing four terminal signals of Udec4x-spec3 probe: one strong on chromosome without Ubriz-spec3 signals, and three weak on chromosomes showing Ubriz-spec3 signals. Scale bars = 5µm

**FIG. 6.**
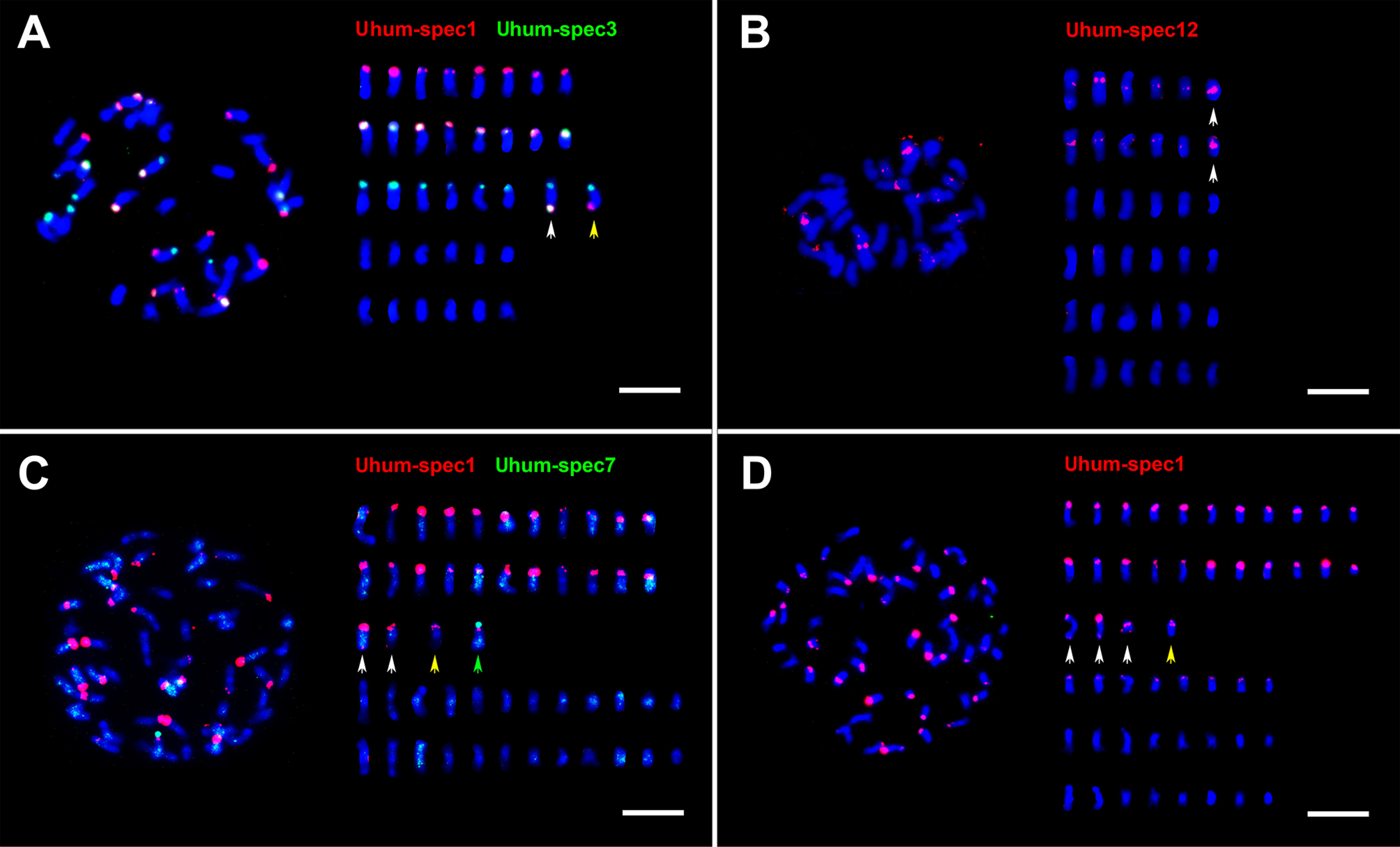
Localization of abundant repetitive sequences on chromosomes of *U. humidicola* accessions. Probes are described in Table S3; Chromosomes (right) were arranged by size and FISH signal. (A) Uhum-spec1 and Uhum-spec3 probes differentiate chromosomes of *U. humidicola* (6*x*, CIAT 26151) into four types: 8 chromosomes with terminal Uhum-spec1 signals (group I), 8 chromosomes with Uhum-spec1 and Uhum-spec3 signals (group II), 8 chromosomes with Uhum-spec3 signals (group III), and 12 chromosomes without any signals (group IV). Two chromosomes belonging to the group III differed from the other six: one of them has two additional signals of Uhum-spec1 and Uhum-spec3 probes (white arrow), while the other has only one additional signal of Uhum-spec1 probe (yellow arrow); (B) *U. humidicola* (6*x*, CIAT 26151) showed signals of Uhum-spec12 probe at centromeric and intercalary position of twelve chromosomes. The intensity and distribution of these signals indicate the presence of six pairs of chromosomes. Especially one pair of shorter chromosomes exhibits very strong centromeric-pericentromeric signals of Uhum-spec12 (white arrows); (C) *U. humidicola* (8*x*+2 or 9*x*-4, CIAT 16867); Uhum-spec7 signals are dispersed along chromosomes, some of them are more intense, but it is difficult to deduce if there is any specific pattern of their distribution (high stringency conditions).Uhum-spec1 probe showed signals on 26 chromosomes, but 4 chromosomes were different, showing additional signals: two chromosomes had extra Uhum-spec1 signals on the opposite arms (white arrows), 1 chromosome showed doubled Uhum-spec1 signal (yellow arrow), and 1 chromosome had strong terminal Uhum-spec7 signal (green arrow); (D) chromosomes of *U. humidicola* (8*x*+2 or 9*x*-4, CIAT 16867); the low stringency conditions, allowing hybridization between DNAs sharing 72% sequence identity, revealed 8 additional weak signals of Uhum-spec1 probe . Three chromosomes have Uhum-spec1 signals on both arms (white arrows). Scale bars = 5µm

**FIG. 7.**
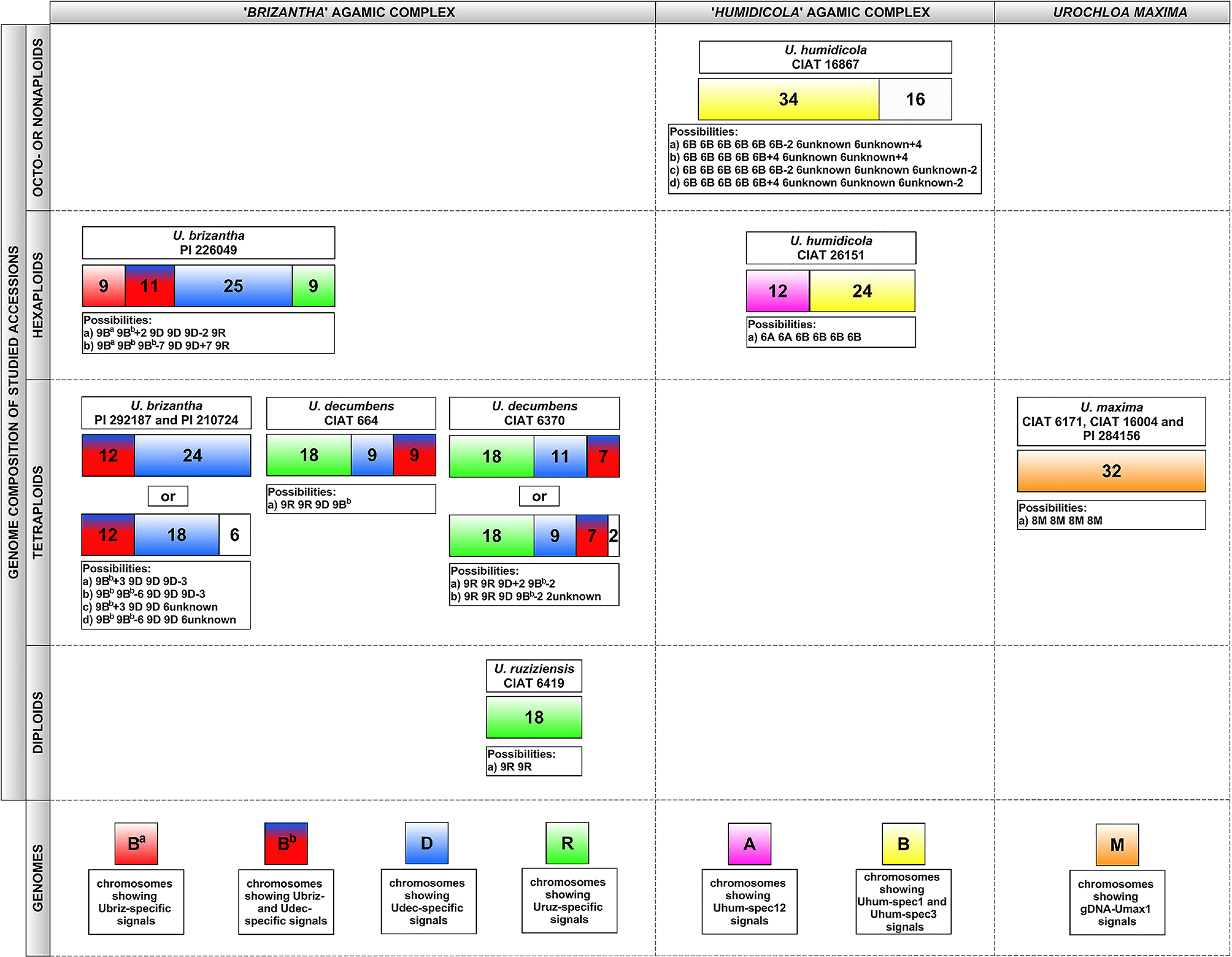
Inferred genomic composition (coloured blocks) and chromosome numbers (shown) of studied accessions belonging to ‘*brizantha*’ (left) and ‘*humidicola*’ (middle) complexes, and *U. maxima* (right). White blocks with numbers: undetermined or diverged genome.

## Discussion

Through our analysis of repetitive DNA sequences using unassembled raw-reads, molecular cytogenetic and flow cytometry tools, we were able to define the nature and similarity between the *Urochloa* species and genomes available internationally in germplasm resource collections (CIAT, USDA, and replicates elsewhere). By identifying repetitive sequences that were unique to the different genomes present in the species, and identifying distinct genomes in the polyploids, we could reveal the genome composition of polyploids and the nature of evolutionary changes in the primary DNA sequence of repetitive motifs and changes in their abundance. Together with growth habit and morphological data, the evaluation of the *Urochloa* material confirmed the challenges in defining the genetic relationships of the accessions used in forage breeding.

### Ploidy and geographical origin

All the species analysed here (**Fig. 1**; **Supplementary Data Table S1**) are native to Africa: *U. brizantha* is widespread in sub-Saharan Africa, the range of *U. decumbens* and *U. ruziziensis* is restricted to the area of Lake Victoria, and *U. humidicola* occurs from Nigeria eastwards to Southern Ethiopia and southwards to South Africa (Renvoize and Maass, 1993). Information about the collection sites, most from the international 1984/85 expeditions representing the majority of germplasm in Colombia and Brazil, and reintroductions within Africa (Wassie *et al*., 2018) allowed us to correlate geographical distribution and ploidy levels as determined by flow cytometry (**Fig. 1A, B**). For *Urochloa* species with multiple ploidies, representatives of all ploidies were found in each geographical region, indicating co-occurrence, no major niche specialization, and opportunity for hybridization and introgression, including segmental allopolyploidy. This is not uncommon for species with multiple ploidies. In wild Asteraceae species *Tripleurospermum inodorum* in central Europe, for example, Čertner *et al*. (2017) studying the spatio-temporal patterns of ploidy coexistence found tetraploid cytotypes alone in about half or more of the populations, diploids in about 10% of populations, with remaining populations being a mixture of ploidies. Natural selection may produce polyploids and hybrids with strong geographical signals (Hagl *et al*., 2021; Alix *et al*., 2017). Even in species with no significant ecological differences between cytotypes (e.g. in *Aster amellus*), no mixing of ploidies is seen even in contact zones (Mandáková and Münzbergová, 2006). Deliberate or accidental roadside or forage introductions (likely to be over-represented in the genebank material sampled here) may introduce different ploidies, although our accessions are genetically different (Hanley *et al*., 2020; Higgins *et al*., in preparation). Polyploids are often argued to have a competitive advantage over diploids (Alix *et al*., 2017) and production of polyploid seeds and individuals by diploids is widespread, although subsequent establishment of whole polyploid populations and their expansion can be hindered by insufficient seed production (Levin, 2021). Thus, it is not surprising that multiple ploidy levels (2x-9x) in many collection areas, including new polyploids and fertile 3x hybrids (**Fig. 1**), were found and suggests co-existence of the various ploidy levels in both *U. brizantha* and *U. humidicola*.

### Chromosome and genome differentiation in Urochloa polyploids

The karyotypes of three *Urochloa* species belonging to ‘*brizantha*’ complex show little differences, having chromosomes similar in size and morphology (Bernini and Marin-Morales, 2001; Nielen *et al*., 2010). Physical mapping of 5S and 18S-5.8S-25S rDNA locations gives a chromosome marker, but mostly similar patterns in the ‘*brizantha*’ complex did not assist in identification of genome composition (**Fig. 2**; Akiyama *et al*., 2010; Nielen *et al*., 2010; Santos *et al*., 2015; Nani *et al*., 2018). In the three accessions with desirable agronomic characteristics which could not be assigned to species based on morphology, the number of rDNA loci did not correspond to ploidy, with only two 45S rDNA loci in the two tetraploids (four sites expected), and four 45S loci in a pentaploid (expectation five), suggesting more complex origin involving processes such as karyotype reorganization, aneuploidy or segmental allopolyploidy and introgression.

Using two diploid total genomic, gDNA, probes (Uruz and Udec) to chromosomes of three species belonging to the ‘*brizantha*’ complex (**Fig. 4A-J**), *in situ* hybridization results showed very small differences in hybridization patterns between groups of chromosomes (candidate genomes), with strong signals in centromeres consistent with Corrêa *et al*. (2020). However, the genome-specific motifs identified in sequence data (see below) suggested some chromosomes sharing similar centromeric signals actually belong to different genomes. Polyploid *U. humidicola* showed dispersed signals of gDNA probes along all chromosomes, making it impossible to discriminate genomes (**Fig. 4K**). Genomic *in situ* hybridization indicated that tetraploid *U. maxima* is autopolyploid (**Fig. 4L**) which is in contrast to the other polypoids in *Urochloa* that have been identified as allopolyploids.

Santos *et al*. (2015) revealed some differentiation of candidate genome-specific Ty3-gypsy retrotransposons in pericentromeric regions of *Urochloa* chromosomes. *Urochloa* contrasts with another Poaceae, *Avena*, where genomic *in situ* hybridization identifies the genomes (Katsiotis *et al*., 2000; Tomaszewska and Kosina, 2021), ‘painting’ most of the chromosomal lengths. The *Urochloa* results indicate that bulk repetitive sequences present in the gDNA probes have diverged only slightly in sequence and copy number during speciation of the diploid *U. brizantha* ancestors combined in polyploids (**Fig. 4**), showing only weak genome-specificity (Corrêa *et al*., 2020). Centromeres of plants are often composed of abundant tandemly repeated sequences and sometimes centromere-specific retrotransposon families (e.g. the CR family in grasses, Miller *et al*., 1998; Presting *et al*., 1998; Heslop-Harrison and Schwarzacher, 2011) Centromere-specific distribution pattern of signals of genomic (**Fig. 4**) and transposable element (Santos *et al*., 2015) probes in *Urochloa* may be due to the retrotransposons being clustered in centromeres and thus generating strong signals, whereas copies located along chromosome arms are dispersed (Miller *et al*., 1998).

While genomic *in situ* hybridization did not differentiate *Urochloa* genomes, bioinformatic analysis of unassembled raw DNA sequence identified short sequence motifs that showed differential abundance among accessions. *In situ* hybridization of the various motifs to metaphase chromosomes confirmed the differential abundance and enabled identification of the genomes present in polyploids (**Figs. 5, 6**), leading to a model of *Urochloa* evolution (see below). All the sequences were present on multiple chromosomes, showing both amplification and dispersion or homogenization of the motifs after speciation from a common ancestral *Urochloa* genome, and each sequence had a characteristic proximal, distal, or more dispersed chromosomal location. However, in contrast to a parallel analysis in *Avena* species (Liu *et al*., 2019), no major DNA satellite or tandem repeats giving chromosomal bands were revealed in *Urochloa*. Triticeae species with much larger genomes and chromosomes have many tandem repeats, including simple sequence motifs, that are tribe, genus or species specific and have been widely used to identify chromosomes (along with total genomic DNA; e.g. Ali *et al*., 2016; Patokar *et al*., 2016). More generally, in a wide range of species, repetitive sequences have been identified as a key component of evolutionary mechanisms and karyotypic differentiation, playing an important role in speciation (Heslop-Harrison and Schwarzacher, 2011; Mehrotra and Goyal, 2014). Comparison of genomic *in situ* hybridization, and the sequences and chromosomal distribution of repetitive sequences identified by cloning or sequence analysis, suggests considerable differences in repetitive sequence evolution between taxonomic ‘groups’ (family, tribe, or genus). It is evident that each group has distinctive rules for chromosome and repetitive sequence evolution, but these are not easily transferrable as models between species groups.

### Taxonomy and the genomic composition of Urochloa polyploids

Species concepts for many of the genebank accessions of *Urochloa* (including *Brachiaria*, and other species which have previously been placed in the genera *Megathyrsus, Eriochloa* and *Panicum*) have been problematic, not least because of the range of ploidies, apomixis, vegetative propagated lines, intermediate morphological traits (**Fig. 1C**; **Supplementary Data Table S1**), growth habits, and the presence of hybrids occurring in the wild or as landraces selected by forage grass breeders and farmers. Our results support the maintenance of distinct species for *U. ruziziensis, U. brizantha, U. decumbens, U. humidicola,* and *U. maxima* (chromosomal organization in **Figs. 4, 5, 6**; relationship models **Figs. 7, 8**).

Following the genome labelling system adopted across the Triticeae (Hordeae) tribe (Linde-Laursen *et al*., 1997) or in *Brassica* (Cheng *et al*., 2013; Alix *et al*., 2008), the level of genomic differentiation as found here by sequence and chromosomal analysis is high enough that we propose designating basic genomes in *Urochloa* using the upper-case letters R, B, and D (for ‘*brizantha*’ complex), A and B or even C (for ‘*humidicola*’ complex), and M for *U. maxima* (**Fig. 7, 8**). More limited differentiation allows us to suggest that superscript designations, referring to modified basic genomes, for less-well differentiated genomes including B^a^ and B^b^. **Fig. 7** illustrates the chromosome and genome composition of the accessions studied here. *U. ruziziensis* was diploid; *U. brizantha* with multiple polyploid levels shows a variation of chromosomes and genomes, as does *U. decumbens*. *U. maxima* contains only the M genome so exists as an auto-tetraploid. Our analysis supported the genome composition of hexaploid *U. humidicola* (based on meiotic behaviour, Vigna *et al*., 2016; and transposable elements, Santos *et al*., 2015) as including A and B genomes (and probably the C genome in higher ploidy levels). Ty1-gypsy Tat probe (Santos *et al*., 2015) and Uhum-spec12 (**Fig. 6B**) are good markers for the A genome. The B genome is more variable in showing three types of chromosomes.

### Evolutionary model for Urochloa species

Three substantive models (**Fig. 8**) to explain evolution of *Urochloa* polyploids in the ‘*brizantha’* and ‘*humidicola’* agamic complexes, and *U. maxima* were generated from multiple lines of evidence. Renvoize and Maass (1993) suggested that diploid *U. decumbens* evolved from *U. brizantha*: the natural range of *U. decumbens* covers the area of a candidate ancestral *U. brizantha* form or variety (e.g. *U. brizantha* var. *latifolium* Oliver or *U. brizantha* var. *angustifolia* Stent & Rattray) with lanceolate hairy leaves and a decumbent habit. We found genome-specific repetitive sequences in *U. decumbens*, but all of them were shared with *U. brizantha,* supporting the order of evolutionary branching. These data contradict Basappa’s *et al*. (1987) suggestion that *U. decumbens* is a natural hybrid between *U. brizantha* and *U. ruziziensis*, and confirmation of this hypothesis would be meiotic abnormalities found in *U. decumbens*. We support this hypothesis for tetraploid *U. decumbens*, but not the diploid accession we studied, although our results were inconclusive for the hexaploid and pentaploid *U. decumbens* accessions (**Fig. 7**). Pessoa-Filho *et al*. (2017) found that tetraploid *U. brizantha* and *U. decumbens* show high similarity of their plastid sequences and low number of SNPs, which may suggest that a single polyploidization event took place to establish both the tetraploid *U. brizantha* and *U. decumbens*: namely a potential fertilization of a tetraploid *U. brizantha* BD gamete and an unreduced RR gamete of a diploid *U. ruziziensis*.

**FIG. 8.**
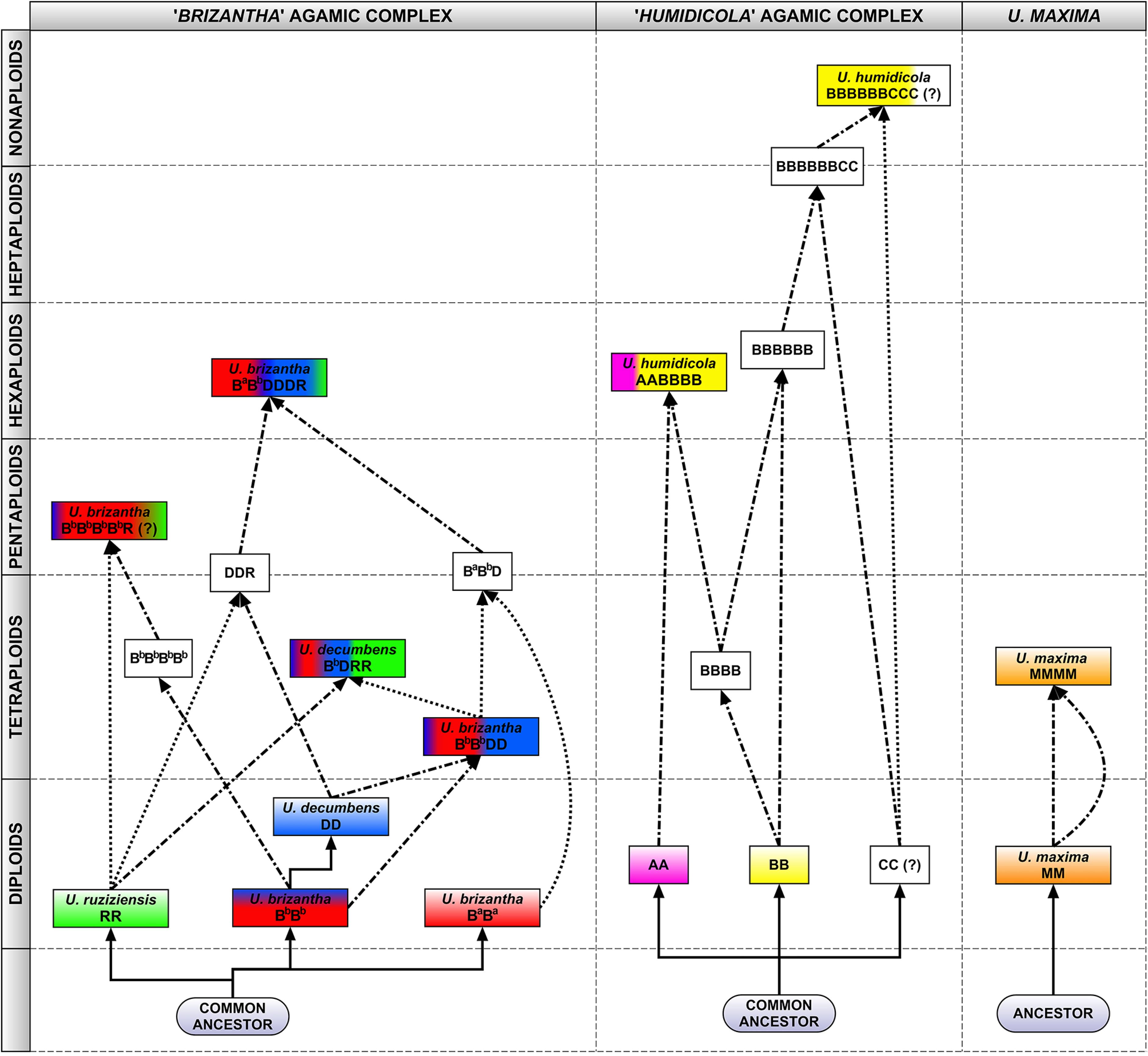
A model for evolutionary origin of *Urochloa* species in ‘*brizantha*’ (left) and ‘*humidicola*’ (middle) complexes, and in *U. maxima* (right), built from this study and published data: genome sizes and ploidy (SUpplementary Data Table S1), repetitive DNA sequences from whole genome sequence analysis (k-mer counts and graph-based clustering; Supplementary Data Tables S3, Supplementary Data Table S9), *in situ* hybridization with defined repeat probes (Figs. 5, 6) and genomic DNA (Fig. 4; and Corrêa *et al*., 2020), karyotype analyses (Corrêa *et al*., 2020), meiotic behaviour (Risso-Pascotto *et al*., 2005; Mendes-Bonato *et al*., 2007; Fuzinatto *et al*., 2007), chloroplast genome (Pessoa-Filho *et al*., 2017), hybrid occurrence (Table 1; Mendes *et al*., 2006; Vigna *et al*., 2016; Risso-Pascotto *et al*., 2005); CIAT breeding programs (Renvoize and Maass, 1993; Miles *et al*., 1996) and reported apomixis (Roche *et al*., 2001). Three line types show evolutionary sequence divergence (solid line), and hybridization events involving haploid, *n*, (dotted line) or unreduced, 2*n*, (dash-dotted line) gametes from different genomes (designated at Fig. 7). White blocks: putative species/hybrids.

Risso-Pascotto *et al*. (2006) suggested that hexaploid *U. brizantha* probably resulted from “chromosome doubling of a triploid derived from species that did not display the same behaviour for spindle organization”. Triploid hybrids were found in nature (Timbó *et al*., 2016), and may originate from crosses between diploid *U. ruziziensis* and tetraploid *U. decumbens* or *U. brizantha*. Thus, a hexaploid species would be created by crossing two different triploids rather than doubling of genomes of a triploid hybrid. This suggestion arises from the presence of only one R genome in the hexaploid *U. brizantha*, as indicated by our *in situ* hybridization analysis (**Figs. 4, 5** and **7**). We also suggest that there are at least two cytotypes/varieties of diploid *U. brizantha* (from *in situ* hybridization and repetitive sequence composition in hexaploid *U. brizantha*), supported by Bernini and Marin-Morales (2001) and Nielen *et al*. (2010) showing differences in karyotypes of diploid *U. brizantha* accessions.

The most likely evolution of species belonging to the ‘*humidicola*’ complex is much more difficult to propose, because all accessions are polyploid and there is no suggestion as to which diploid species may be considered ancestral. There are three known levels of ploidy in this species: hexaploid, heptaploid and nonaploid (Boldrini *et al*., 2009a; Jungmann *et al*., 2010; Vigna *et al*., 2016; we also had an inconclusive 2*n*=50 accession). Our analysis of the genomic composition of the hexaploid species coincides with meiotic analyses conducted by Boldrini *et al*. (2009b) and Vigna *et al*. (2016), and the model of evolution of species belonging to ‘*humidicola*’ complex is supported by *in situ* hybridization with specific probes (**Fig. 6**). The B genome includes chromosomes showing three different types of signals, which may suggest that *U. humidicola* has gone through several rounds of polyploidization. Analysis of the genomic composition of *U. dictyoneura*, which belongs to the ‘*humidicola*’ complex would be desirable, especially since tetraploid accessions with 2*n*=4*x*=24 are known (Boldrini *et al*., 2010) and could have contributed to the evolution of *U. humidicola*, which shows odd ploidy levels.

Our *in situ* hybridization studies gave evidence for potential introgression within *Urochloa.* Some polyploid lines (*U. brizantha* and *U. humidicola*) here have chromosome pairs that are different from others within their genome (**Figs 5G, J, K, L**; **6A, C, D**), looking like segmental allopolyploidy (Mendes-Bonato *et al*., 2002) or disomic introgression lines. Frequent introgression seems to occur in wheat (Cheng *et al*., 2019) and oat polyploids, and in breeding, whole chromosomes, chromosome arms or segments, may be substituted. An example is *Triticale* which may have not the expected 7 chromosome pairs of each genome but 14 A, 12 B, 2 D and 14 R chromosomes (Neves *et al*., 1997). Some hybrid species are diploid or reduce chromosome numbers so they are not clearly tetraploid – *Petunia hybrida* is 2*n*=14, like its ancestors (Bombarely *et al*., 2016), with mixture of ancestral genomes, while the octaploid *Nicotiana* cell fusion hybrid (4*x* + 4*x*) has lost a few chromosomes (Patel *et al*., 2011).

### Conclusions

Genome composition and evolution are complex in *Urochola* tropical forage grasses. Grasslands are not only a major source of food production but also environmental services: water, soil preservation, carbon capture etc., often in more biodiverse regions, where identification of species and their relationships will assist grass conservation. Despite their lower economic value, breeding and exploitation of biodiversity is required within the group (whether using sequence data or a genetic map, as in e.g. *Lolium*, Tomaszewski *et al*., 2012). Like wheat and *Brassica* crops, wild relatives contribute to the current pool of diversity used in *Urochloa* tropical forage grass improvement, with additional complexities from apomixis. The knowledge of genome relationships and polyploid genome composition gives opportunities for rational and systematic use the accessions in forage improvement programmes (superdomestication: Vaughan *et al*., 2007). Complementing our study showing the diversification of genomes and repetitive DNA, a parallel study (Hanley *et al*., 2020) found high levels of genetic diversity in 20 genes related to forage quality in 104 of the accessions studied here.

Our study was focused on accessions available from international germplasm collections to breeders and researchers. As Keller-Grein *et al*. (1996) correctly pointed out, further collecting of the *Urochloa* species in Africa would be worthwhile to enrich the germplasm collection with new accessions, finding further useful characteristics which can be exploited, and to better understand complicated evolution, adding to the analysis here. For legal regulations regarding biosecurity restrictions (diseases and invasive species) and Plant Breeders Rights and germplasm ownership, it is necessary to have an accepted name for every species, and our identification of genomes and genome composition in *Urochloa* polyploids presents the necessary framework.

Breeding programs often work with a single ploidy as directed crosses among parents with different ploidies are challenging. We would suggest that *Urochloa* species are all part of a common gene pool, and any hybrid combination might be possible and become a successful forage variety, noxious weed or disease host. The current breeding programs at CIAT manage tetraploid interspecific crosses within the ‘*brizantha*’ agamic complex, hexaploid crosses within the ‘*humidicola*’ agamic complex, and tetraploid intraspecific crosses of *U. maxima*. Choice of appropriate strategies to generate hybrids requires knowledge of ploidy provided by our research, supported by the model of evolution and diversification of the species.

## Acknowledgements

This work was supported under the RCUK-CIAT Newton-Caldas Initiative “Exploiting biodiversity in *Brachiaria* and *Panicum* tropical forage grasses using genetics to improve livelihoods and sustainability”, with funding from UK’s Official Development Assistance Newton Fund awarded by UK Biotechnology and Biological Sciences Research Council (BB/R022828/1). PT has received further support (polyploidy and chromosome evolution) from the European Union’s Horizon 2020 research and innovation programme under the Marie Sklodowska-Curie grant agreements No 844564 and No 101006417.

We would like to thank Dr Jennifer Hincks from The Centre for Core Biotechnology Services, Flow Cytometry Facility at University of Leicester, for assistance with developing our flow cytometry protocol. We are grateful to USDA-Germplasm Resources Information Network (GRIN) for their generous provision of seeds. Germplasm held in the CIAT collections is available on request (http://genebank.ciat.cgiar.org).

## Additional files

**Supplementary Data Fig. S1** Ploidy measured by flow cytometry of PI-stained nuclei from dehydrated tissues of diploid (2*x*), tetraploid (4*x*), pentaploid (5*x*), and hexaploid (6*x*) accessions of *Urochloa* showing very sharp peaks (measured by CV, coefficient of variation). Regions of identification (red) were placed across the peaks to export values representing peak positions and CVs.

**Supplementary Data Fig. S2** Contig 5 as a candidate motif specific to *U. brizantha* genome.

**Supplementary Data Fig. S3** Distribution of graph-based clusters. Hierarchical agglomeration of RepeatExplorer and TAREAN analyses of *Urochloa* accessions genomes are shown. X-axis denotes the cumulative read number percentage while Y-axis denotes the read numbers in the clusters.

**Supplementary Data Table S1** List of accessions used in the study, their ploidy levels, growth habitats and geographical distribution.

**Supplementary Data Table S2** Summary of sequencing data quality.

**Supplementary Data Table S3** Potential genome-specific 50-mer sequences, their genome proportion, and description of probes and *in situ* hybridization signals.

**Supplementary Data Table S4** BLASTN search of highly abundant potential genome-specific 50-mers.

**Supplementary Data Table S5** RepeatExplorer characterization of selected repeat clusters of *Urochloa* accessions.

**Supplementary Data Table S6** The NCBI BLASTN results of clusters found using RepeatExplorer. Description, query coverage and identity are listed for each cluster.

**Supplementary Data Table S7** TAREAN characterization of selected repeat clusters of *Urochloa* accessions.

**Supplementary Data Table S8** The NCBI BLASTN results of clusters found using TAREAN. Description, query coverage and identity are listed for each cluster.

**Supplementary Data Table S9** Potential genome specific repeats and their genome proportion. Description of *in situ* hybridization results and information about promising genome-specific sequences not tested as probes are given.

**Supplementary Data Table S10** Repetitive DNA composition of *Urochloa* genomes. Genome portion (as percent) is listed for each species. Families and subfamilies of Transposable elements Class I (retrotransposons) and Class II (DNA transposons), and Tandem repeats are also given.

**Supplementary Data Table S11** List of genomic DNA probes.

**Supplementary_Data _Table_S12** Genomic *in situ* hybridization results.

## Notes

### Competing Interest Statement

The authors have declared no competing interest.

