## Supplementary figures and images for "Complex polyploid and hybrid species in an apomictic and sexual tropical forage grass group: genomic composition and evolution in *Urochloa* (*Brachiaria*) species"

### Supplementary Data Fig._S1

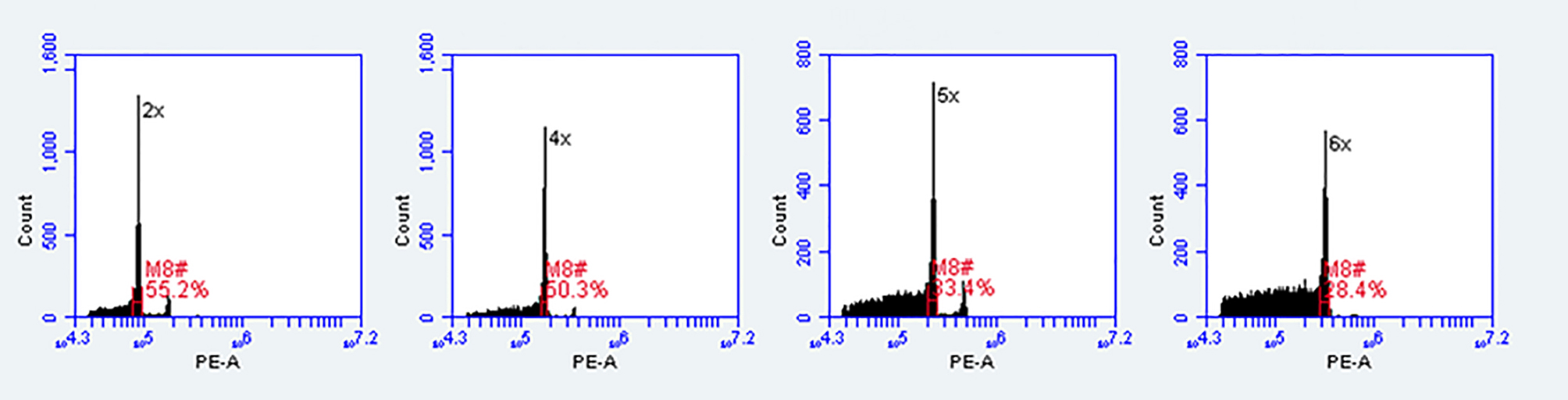

### Supplementary Data Fig._S3

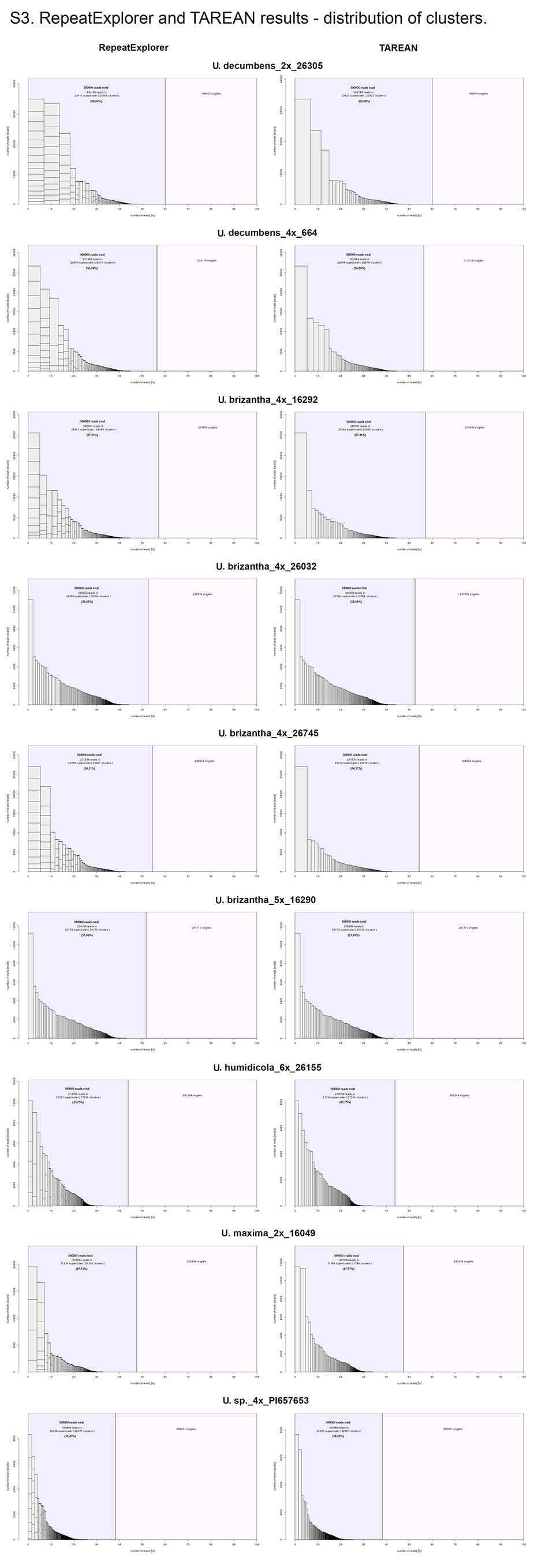
