## Supplementary Data Fig._S2 for "Complex polyploid and hybrid species in an apomictic and sexual tropical forage grass group: genomic composition and evolution in *Urochloa* (*Brachiaria*) species"

**Fig. S2. Contig 5 as a candidate motif specific to *U. brizantha* genome**

**Contig5_U.brizantha** AGAATGTTGTCTGCTACACAGGGCGAGGCAAATTTACATGAATAGAGCTATCATGATTTTTTAGAAAAAAA=TAATAACTCCCACATGTCCATAGAGGTACCCCTTGGGCATGCAGCGTCGCAGCTCACGAGAGGGGCTCCAGCGCACTACCATGGATGGTCCACATCTCAAGTAGTAGTCGAGGAACACCAAGACATGGTATGTTAACTCATATGTTAATTTCGGGTTGTGGACCAAACGTCTAGTCCATAGTCTTGGCGTCCGAGTGGGTTTCGACGGTCAAAATTGATTAACTTCCCCATGAATTTCCATAACTTATTCGTTTGGAGCCCGAATCAATCGCGTTTTTT

Length: 350 bp & 1 gap

GC: 44.6%

Genome proportion:

|  | *U. brizantha*  CIAT 26745 (4*x*) | *U. brizantha*  CIAT 26032 (4*x*) | *U. brizantha*  CIAT 16292 (4*x*) | *U. brizantha*  CIAT 26745 (5*x*) |
| --- | --- | --- | --- | --- |
| Contig 5 | 1,03% | 0,26% | 0,33% | 0,92% |
