## Supplementary Data Table_S2 for "Complex polyploid and hybrid species in an apomictic and sexual tropical forage grass group: genomic composition and evolution in *Urochloa* (*Brachiaria*) species"

###### Table S2. Summary of sequencing data quality.

| Species | Accession number | Donor | No. of chromosomes | Raw reads | Raw data (G) | Effective (%) | Error (%) | Q20 (%) | Q30 (%) | GC (%) |
| --- | --- | --- | --- | --- | --- | --- | --- | --- | --- | --- |
| *Urochloa* sp. | PI 657653 | USDA | 2*n*=4*x*=32 | 49413044 | 14.8 | 99.50 | 0.02 | 94.67 | 88.57 | 45.55 |
| *U. brizantha* | 16290 | CIAT | 2*n*=5*x*=45 | 33442671 | 10.0 | 99.32 | 0.03 | 94.30 | 87.05 | 45.84 |
| *U. brizantha* | 26032 | CIAT | 2*n*=4*x*=36 | 33712395 | 10.1 | 99.25 | 0.02 | 94.72 | 87.78 | 45.92 |
| *U. brizantha* | 26745 | CIAT | 2*n*=4*x*=36 | 39939853 | 12.0 | 99.31 | 0.01 | 95.80 | 90.52 | 45.08 |
| *U. brizantha* | 16292 | CIAT | 2*n*=4*x*=36 | 50993234 | 15.3 | 98.92 | 0.02 | 95.16 | 89.21 | 47.04 |
| *U. decumbens* | 664 | CIAT | 2*n*=4*x*=36 | 33536782 | 10.1 | 99.03 | 0.01 | 96.70 | 92.03 | 44.91 |
| *U. decumbens* | 26305 | CIAT | 2*n*=2*x*=18 | 39753626 | 11.9 | 99.50 | 0.01 | 96.40 | 91.82 | 46.67 |
| *U. humidicola* | 26155 | CIAT | 2*n*=6*x*=36 | 42095385 | 12.6 | 98.66 | 0.02 | 94.57 | 88.50 | 44.75 |
| *U. maxima* | 16049 | CIAT | 2*n*=2*x*=16 | 42391844 | 12.7 | 98.55 | 0.02 | 94.64 | 88.35 | 45.60 |

Raw reads: four rows are taken as a unit to calculate the total amount of read1 and read2 in raw data files; Raw bases: (total raw reads) * (sequence length), calculating in G; Error rate: base error rate; Q20, Q30: (Base count of Phred value > 20 or 30) / (Total base count); GC content: (G & C base count) / (Total base count)
