## Supplementary Data Table_S11 for "Complex polyploid and hybrid species in an apomictic and sexual tropical forage grass group: genomic composition and evolution in *Urochloa* (*Brachiaria*) species"

**Table S11.** Whole genomic DNA used as probes for *in situ* hybridization.

| **Species** | **Accession number** | **Donor** | **Name of probe** | **Number of chromosomes** |
| --- | --- | --- | --- | --- |
| *U. brizantha* | 16341 | CIAT | gDNA_Ubriz1 | 2*n*=2*x*=18 |
| *U. decumbens* | 26305 | CIAT | gDNA_Udec1 | 2*n*=2*x*=18 |
| *U. decumbens* | 6133a | CIAT | gDNA_Udec2 | 2*n*=2*x*=18 |
| *U. ruziziensis* | 26348 | CIAT | gDNA_Uruz1 | 2*n*=2*x*=18 |
| *U. maxima* | 16049 | CIAT | gDNA_Umax1 | 2*n*=2*x*=16 |
| *U. maxima* | 6898 | CIAT | gDNA_Umax2 | 2*n*=2*x*=16 |
