## Supplementary Data Table_S12 for "Complex polyploid and hybrid species in an apomictic and sexual tropical forage grass group: genomic composition and evolution in *Urochloa* (*Brachiaria*) species"

**Table S12.** Genomic *in situ* hybridization results.

| Species | Accession number | Genomic DNA probes | | | |
| --- | --- | --- | --- | --- | --- |
|  |  | Ubriz | Udec | Uruz | Umax |
| *U. ruziziensis* | CIAT 6419 | no signal | 18 chromosomes showing signals in centromeres | 18 chromosomes showing signals in centromeres; same position of signals as Udec probe | no signal |
| *U. decumbens* | PI 210724 | 4 rDNA signals | 18 chromosomes show strong signals, 18 chromosomes with weak signals in centromeres | 18 chromosomes show strong signals, 18 chromosomes with weak signals in centromeres; same position of signals as Udec probe | 4 rDNA signals |
| *U. decumbens* | CIAT 664 | 4 rDNA signals | 18 chromosomes show strong signals, 18 chromosomes with weak signals in centromeres | 18 chromosomes show strong signals, 18 chromosomes with weak signals in centromeres; same position of signals as Udec probe | 4 rDNA signals |
| *U. decumbens* | CIAT 6370 | 4 rDNA signals | 18 chromosomes show strong signals, 18 chromosomes with weak signals in centromeres | 18 chromosomes show strong signals, 18 chromosomes with weak signals in centromeres; same position of signals as Udec probe | 4 rDNA signals |
| *U. brizantha* | PI 292187 | 4 rDNA signals | 9 chromosomes painted; 27 chromosomes show centromeric signals | 9 chromosomes painted; 27 chromosomes show centromeric signals; same position of signals as Udec probe | 4 rDNA signals |
| *U. brizantha* | PI 226049 | 6rDNA signals | Signals in centromeres: 18 strong, 18 weaker, 18 weak | Signals in centromeres: 18 strong, 18 weaker, 18 weak | 6 rDNA signals |
| *U. humidicola* | CIAT 26151 | 6 rDNA signals | Interspersed signals along chromosomes | Interspersed signals along chromosomes | 6 rDNA signals |
| *U. humidicola* | CIAT 16867 | 6 rDNA signals | Interspersed signals along chromosomes | Interspersed signals along chromosomes | 6 rDNA signals |
| *U. maxima* | CIAT 6171 | 4 rDNA signals | 4 rDNA signals | 4 rDNA signals | 32 chromosomes show telomeric/ subtelomeric and centromeric signals |
| *U. maxima* | CIAT 16004 | 4 rDNA signals | 4 rDNA signals | 4 rDNA signals | 32 chromosomes show telomeric/ subtelomeric and centromeric signals |
| *U. maxima* | PI 284156 | 4 rDNA signals | 4 rDNA signals | 4 rDNA signals | 32 chromosomes show telomeric/ subtelomeric and centromeric signals |
| *U.* sp. | PI 657653 | 4 rDNA signals, weak signals in centromeres | 4 rDNA signals | 4 rDNA signals | 16 chromosomes painted |
| *U.* sp. | PI 508571 | No signal | 18 chromosomes show signals in centromeres | 18 chromosomes show signals in centromeres; same position as Udec probe | No signal |
| *U.* sp. | PI 508570 | 18 chromosomes painted; some chromosomes show Ubriz and Umax signals | 4 rDNA signals | No signal | 27 (?) chromosomes painted |
